# SYS-Mut: Decoding the Functional Significance of Rare Somatic Mutations in Cancer

**DOI:** 10.1101/2021.09.03.458896

**Authors:** Sirvan Khalighi, Peronne Joseph, Deepak Babu, Salendra Singh, Thomas LaFramboise, Kishore Guda, Vinay Varadan

**Affiliations:** Division of General Medical Sciences-Oncology, Case Comprehensive Cancer Center, Case Western Reserve University, Cleveland, OH-44106 U.S.A; Department of Genetics and genome Sciences, Western Reserve University School of Medicine, Cleveland, OH-44106 U.S.A; Digestive Health Research Institute, Case Western Reserve University School of Medicine, Cleveland, OH-44106 U.S.A; Department of Pathology, Case Western Reserve University School of Medicine, Cleveland, OH-44106 U.S.A

**Keywords:** Multi-omic integrative analysis, glucocorticoid receptor, drug sensitivity, lipid metabolism, functional genomics

## Abstract

Current tailored-therapy efforts in cancer are largely focused on a small number of highly recurrently-mutated driver genes but therapeutic targeting of these oncogenes remains challenging. On the other hand, the vast number of genes mutated infrequently across cancers have received less attention, in part, due to a lack of understanding of their biologic significance. Here we present SYS-Mut, a systems biology platform that can robustly infer the biologic consequences of somatic mutations by integrating routine multi-omic profiles in primary tumors. We established the accuracy of SYS-Mut by recapitulating the functional impact of known driver genes in PanCancer datasets. Subsequent application of SYS-Mut on low-frequency gene mutations in Head and Neck Cancers (HNSC), followed by molecular and pharmacogenetic validation, revealed the lipidogenic network as a novel therapeutic vulnerability in aggressive HNSC. SYS-Mut is thus a robust scalable framework that enables discovery of new targetable avenues in cancer.

## Background

Large-scale profiling studies of multiple cancers have revealed a substantial number of molecular aberrations that accumulate across epigenomic, genomic and transcriptional scales [1,2]. The resulting molecular heterogeneity across cancer lesions within the same tissue type reflect the multiple ways in which signaling networks can be pathologically altered. However, of the plethora of somatic mutations typically observed in a given cancer sample, only a minority are expected to functionally contribute to cancer etiology and/or aggressiveness [2,3]. This molecular heterogeneity coupled with the complexity of co-occurring molecular alterations in individual tumor samples jointly pose a significant challenge to computationally deciphering functionally significant mutations.

Several computational approaches have been developed to analyze cancer mutational profiles to identify and prioritize driver genes and/or pathways [4]. While gene-Level [2,5–8] and network-level [9–12] frequentist approaches have been able to identify the most recurrently mutated genes and pathways associated with cancer initiation and development, they tend to be less reliable in identifying functionally significant genes that fall within the long tail of rarely mutated genes in cancer owing to the sheer diversity of genetic and epigenetic changes in tumors and the significant heterogeneity between and within tumor samples. Furthermore, these frequentist methods suffer from poor specificity because they do not fully account for other molecular measurements and confounding effects of co-occurring pathway deregulations in the network neighborhood while assessing mutation impact. As such, computational approaches that integrate genomic, transcriptomic, and epigenetic data with gene regulatory networks have been developed to provide insight into the impact of somatic mutations on biologic pathways [13–19]. The state-of-the-art approaches have identified a relatively small number of significantly mutated cancer driver genes. However, the functional significance of the vast majority of genes that are mutated at low frequencies (<5%) remains largely unknown (**Supplementary Fig. 1**). One of the key limitations of existing integrative approaches is that they do not explicitly account for the influence of co-occurring *cis*-/*trans*-regulatory factors that could confound the estimates of a given mutation’s downstream transcriptional impact. Indeed, the multi-scale molecular complexity of cancer necessitates the development of integrative analytic frameworks that decode the functional impact of mutational events while accounting for co-occurring *cis*-/*trans*-regulatory molecular aberrations in each tumor sample.

Recognizing the multi-factorial molecular deregulations emblematic of cancer tissues, we present a *SYS*tems biology framework for identification and prioritization of *Mutations* (SYS-Mut) with significant functional impact. SYS-Mut integrates tumor mutational profiles with DNA copy number, DNA Methylation and transcriptomic data to reliably decode the downstream transcriptional impact of rare somatic mutations. We first motivate the theoretical development of the SYS-Mut algorithm based on a hierarchical Bayesian regression framework, followed by establishing the robustness of SYS-Mut through computational benchmarking on simulated data that closely mimics actual PanCancer molecular profiling data. We then showcase the utility of SYS-Mut analyzing multi-omics data derived from the cancer genome atlas (TCGA) [20] dataset, spanning more than 8,000 tumor samples and 29 solid tumor types, identifying both well-known driver genes as well as rare mutational events that modulate key downstream transcriptional programs. We orthogonally assess the transcriptional impact of mutations identified by SYS-Mut using functional genomics assessments in cell line models. Finally, we highlight the translational utility of the SYS-Mut framework by identifying a subnetwork of rarely mutated genes involved in lipid metabolism that are associated with poorer clinical outcomes in Head and Neck cancer and differential sensitivity to drugs targeting the lipid metabolism pathway.

## Methods

SYS-Mut is designed to decode the impact of somatic mutations in a given gene on its downstream transcriptional targets after accounting for likely confounding *cis*- and *trans*-regulatory factors that influence the expression of these target genes. Accordingly, SYS-Mut is structured as a framework of interoperating modules that systematically integrate genome-scale multi-omics profiles of cancer tissues with gene regulatory interaction data to identify mutated genes that significantly impact downstream transcriptional programs (**Fig. 1**; See **Supplementary Information**).

**Fig. 1.**
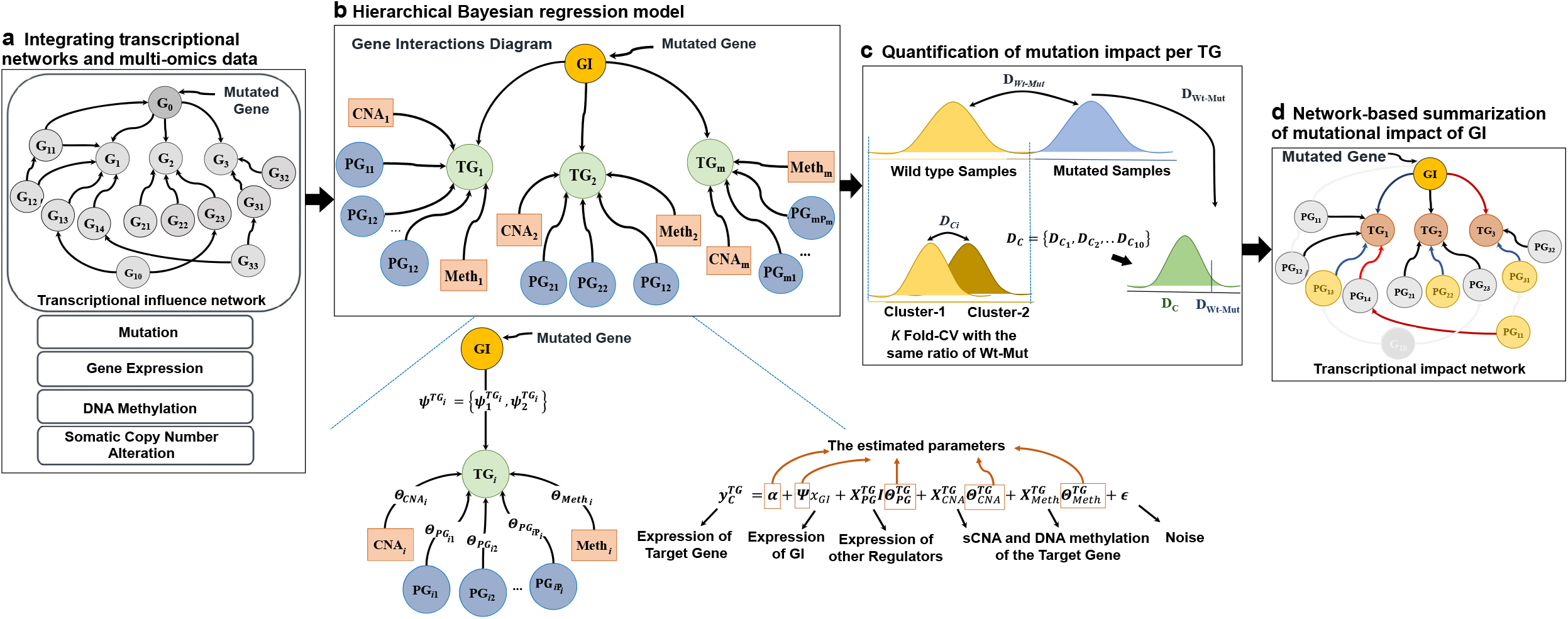
Overview of SYS-Mut algorithmic framework. **a** *Module 1*: For each mutated Gene (G_0_), this module identifies its transcriptional target genes by traversing a predefined transcriptional influence network. Additionally, this module prepares multi-omics profiles of tumor samples for downstream analysis, including mutation status of the gene of interest, gene expression levels of all genes in the network, along with DNA Methylation and somatic copy number alterations (sCNA) of all target genes. **b** *Module 2*: This module employs a hierarchical Bayesian regression framework that integrates multi-omics tumor profiles to estimate the relative influence of each gene of interest (*GI*) on the expression levels of a specific transcriptional target gene (*TG*) in both mutated and non-mutated tumor samples, while also accounting for other *cis*-/*trans*-regulatory influences on the *TG* expression. **c** *Module 3*: This module uses the output of the Bayesian regression framework to determine the statistical significance of differences in the regulatory influence of the *GI* on the expression levels of a specific *TG* in the mutated and non-mutated contexts, thus enabling quantification of the transcriptional impact of mutations in *GI* on the expression levels of *TG*. **d** *Module 4*: Predicts probabilities of the mutational impact of each *GI* by summarizing the impact of *GI* across all downstream *TG*s.

### Module 1: Integrating gene regulatory roadmaps with multi-omics measurements

We first build a genome-scale transcriptional influence network that captures the regulatory roadmaps underlying gene transcription by integrating signaling networks and transcription factor network information from multiple publicly available databases (See **Supplementary Information; Supplementary Table S1**). The resulting directed transcriptional influence network of SYS-Mut includes direct transcriptional targets in addition to close indirect transcriptional regulation via *Complex formation, Abstracts, Families* and/or a single post-translational modification event, resulting in a total of 120655 transcriptional influences spanning 18,651 unique nodes. Next, for each mutated gene, which henceforth is denoted as a gene-of-interest (*GI*), *Module 1* (**Fig. 1a**) traverses the directed transcriptional influence network to identify all of the downstream transcriptional targets (*TG*) potentially impacted by *GI*, while also capturing all of the known *trans*-regulators of each of the *TG*s, denoted henceforth as parent genes (*PG*) (**Supplementary Fig. 2**). In addition, *Module 1* also incorporates known *cis*-regulatory influences, including DNA Methylation (DNAMeth) and somatic copy-number alterations (sCNA) overlapping the gene’s coding and/or promoter regions. *Module 1* then independently processes and normalizes each of the multi-omics data streams of RNA Sequencing (RNASeq), DNAMeth and sCNA along with genome-scale somatic mutation calls to enable downstream integrative analyses.

### Module 2: Hierarchical Bayesian regression model

SYS-Mut *Module 2* (**Fig. 1b**) learns multiple hierarchical regression functions that model the expression levels of the set of transcriptional target genes (*TG*) by accounting for both their *cis*-regulatory factors (*i.e*. DNAMeth and sCNA status) as well as their *trans*-regulatory factors (*i.e*. gene expression levels of all upstream regulators including the mutated *GI*). SYS-Mut model deploys cancer samples that harbor non-synonymous mutations (*Mut*) (see **Supplementary Information**) in a *GI* with samples that are wild-type (*Wt*) for that *GI* and all its connected genes including *TG*s and other trans-regulators (*PG*s) to learn a hierarchical Bayesian regression function equation (1) that models the expression level of the *GI*-associated *TG*s using as input both the *trans*-regulators (mRNA expression of *GI* and other *trans*-regulators, *PG*), as well as *cis*-regulators (sCNA and DNAMeth) of each *TG*:

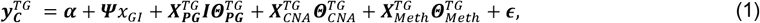

where 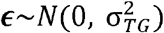.

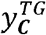 denotes the gene expression levels of the transcriptional target gene *TG*, and ***C*** = {*Wt, Mut*}, corresponds to the wild-type (*Wt*) and mutated (*Mut*) clusters. For the sake of simplicity, the mutated cluster consists of samples harboring non-synonymous mutations in the corresponding *GI*, but in a general case, ***C*** = {*c*_1_, *c*_2_, …, *c_D_*} denotes a vector of *D* clusters that can include a variety of mutation categories such as wild-type, synonymous, missense, nonsense and frame-shift mutations.

*x_GI_* denotes the expression level of a gene of interest *GI*.

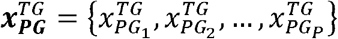 denotes expression levels of the other upstream trans-regulator genes ***PG*** of target gene *TG* except *GI*.

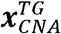 and 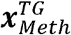 denote *sCNA* and DNAMeth data of the target gene ***TG***, and 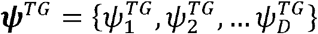 denotes a set of cluster specific parameter 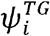 of cluster *i* of the *TG*. These parameters represent mutation effect *GI* on each *TG*. For the case of two clusters, namely wild-type (*Wt*) versus mutated (*Mut*), 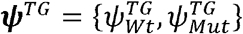 denotes posterior averages of the wild-type and mutated cluster of the *TG*.

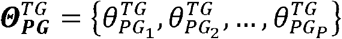 denotes a vector of the estimated parameters of the other upstream trans-regulator genes ***PG*** of target gene *TG* except *GI*.

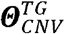 and 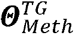 denote the estimated parameters for sCNA and DNAMeth data of each target gene *TG*, respectively, and ***α*** = {α_1_, α_2_, …, α_*D*_}, is vector of cluster specific intercepts.

The objective is to estimate the optimal intra-cluster parameters ***Ψ^TG^***, and inter-cluster parameters 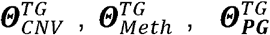 and provide the best prediction power via minimizing the mean squared error between the predicted and observed expression levels of the target gene *TG*. Of note, the parameter inference is marginalized over the cluster-specific parameters **Ψ** (see **Supplementary Information**).

### Module 3: Quantification of mutation impact per *TG*

In order to determine the modulatory role of mutated *GI* on the expression of the downstream *TGs*, the above regression model is applied to n_1_ *Wt* and n_2_ *Mut* samples (**Fig. 1c, Supplementary Fig. 3a**). SYS-Mut provides a function bank, where each function corresponds to a specific *TG* and this function bank represents all *cis*-/*trans*-regulatory alterations. In fact, the prediction process provides the conditional posterior distribution for the expression level of the *TG*. The maximum a-posteriori (MAP) method is used to obtain the expected gene expression levels and parameter values.

As depicted in **Supplementary Figs. 3a and b**, the hierarchical Bayesian regression model estimates inter-cluster parameters 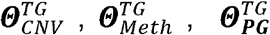 and intra-cluster parameters 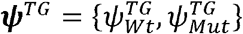, where the difference between the posterior distribution of the cluster specific parameter 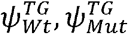 represent the impact of *GI* to the specific *TG*. The Bhattacharyya distance 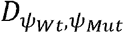between the two posterior distributions 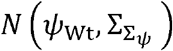 and 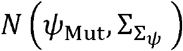 represents the impact of mutations in *GI* on its regulatory influence on the downstream *TG*, with higher *D* corresponding to larger impact (**Supplementary Fig. 3c**).

Given that the parameter estimates and the resulting Bhattacharyya distance, *D_wt–Mut_* may be sensitive to measurement noise and other technical artifacts related to the molecular profiling platforms employed in a given cohort of samples, SYS-Mut next defines a null distribution of *D*, as a control, using the samples that do not harbor any mutations in *GI*. Specifically, as shown in **Supplementary Fig. 3b**, SYS-Mut identifies tumor samples that are wild-type for *GI* as well as all *TG*s of the mutated *GI* and their related parents, *PG*. These ‘*clean wild-type*’ samples are then randomly split into two sub-clusters whose relative sizes are proportional to the sizes of the number of wild-type (*Wt*) and mutated (Mut) samples employed during the estimation of *D_wt–Mut_*. SYS-Mut, then repeats the estimation of cluster specific parameters *ψ*_clust_1_ and *ψ*_clust_2_ thus calculating the Bhattacharyya distance corresponding to a no-impact scenario. This process is then repeated for *K* multiple random splits of the wild-type samples thus generating a null distribution of Bhattacharyya distances ***D***_*c*_ = {*D*_*c*_1__, *D*_c_2__, …, *D_C_k__*}. SYS-Mut then assesses whether *D_wt–Mut_* is significantly greater than ***D***_*c*_ (**Fig. 1c, and Supplementary Fig. 3b, c**) thus determining the statistical significance of the transcriptional impact of the mutated *GI* on the specific *TG*.

### Module 4: Network based summarization of mutational impact of *GI*

To determine the level of mutational impact of a *GI* on each of its downstream *TGs*, SYS-Mut, *module 4* (**Fig. 1d**), next assesses the statistical significance of transcriptional impact using a network randomization strategy. After assessing the log likelihood of impact status for each transcriptional targets (*TG*) of a mutated *GI*, SYS-Mut then replaces the target gene and its upstream *trans*-regulatory network with a random *TG* and its associated *trans*-regulatory network. SYS-Mut then assess the log likelihood of impact status for a total *N* independent random trials of the replaced *TGs*, aiming to generate a *null distribution* of impact. The statistical difference in a *TG’s* log likelihood of impact and the *null distribution* of log likelihoods of random *TGs* indicates the impact score, corresponding to the mutation impact level of a *GI* to its downstream *TG*.

Additionally, to determine the overall impact of mutations in a *GI*, SYS-Mut, *module 4* (**Fig. 1d**), summarizes the mutational impact of the *GI* across all its *TG*s by estimating the fraction of *TG*s whose transcriptional patterns were significantly impacted by mutations in *GI*. SYS-Mut additionally generates a *null distribution* of the fraction of random *TGs* impacted by mutations in a given *GI*. *GI*s whose observed fraction of target genes impacted were significantly (*P* < 0.05) higher than the *null distribution* were deemed to be functionally significant (**Supplementary Fig. 4**). The *GI*-level impact score, which is the average of *GI* to *TGs* impact scores, is used to rank the mutated genes, where higher scores are correspond to higher mutational impact.

## Results

### Computational benchmarking of SYS-Mut

In order to assess the effectiveness of SYS-Mut in identifying mutations with significant functional impact, we performed extensive computational benchmarking experiments using simulated data that mimic real-world multi-omic profiles of cancer. Accordingly, we applied SYS-Mut on a large set of simulated mutation, gene expression, sCNA and DNAMeth data generated using the model described in equation (1). Taken together, we performed simulation-based benchmarking of SYS-Mut’s performance to detect low, medium or high impact of mutations in GI on downstream transcriptional target gene (*TG*) expression. The simulation studies were designed to evaluate SYS-Mut performance across a variety of conditions including, number of samples (100, 300, 500, and 700) available for training the model, mutation rates of GI (0.5%, 1%, 2%, and 5%), and number of *trans*-regulators of TG (5, 10, 50, 100) that could confound the impact mutation assessment. The AUC curves related to the aforementioned simulation experiments (**Figs. 2a, b, c**), confirm the model’s ability to identify rare mutations with significant downstream impact even when the sample size is low (*N*=100), the mutation rate of the GI is low (<1%) and the number of potential *trans*-regulatory confounders for a given TG is high (>100). Importantly, these simulation studies also reveal the importance of SYS-Mut’s approach to explicitly account for *cis-/trans*-regulatory factors that modulate TG expression levels, thus ensuring robust detection of the downstream transcriptional impact of mutations in *GI* (**Fig. 2c**). Our simulation studies, therefore, reveal a key advantage of SYS-Mut as compared to algorithms such as Xseq[15] whose estimates are liable to be confounded by such *cis-/trans*-regulatory factors.

**Fig. 2.**
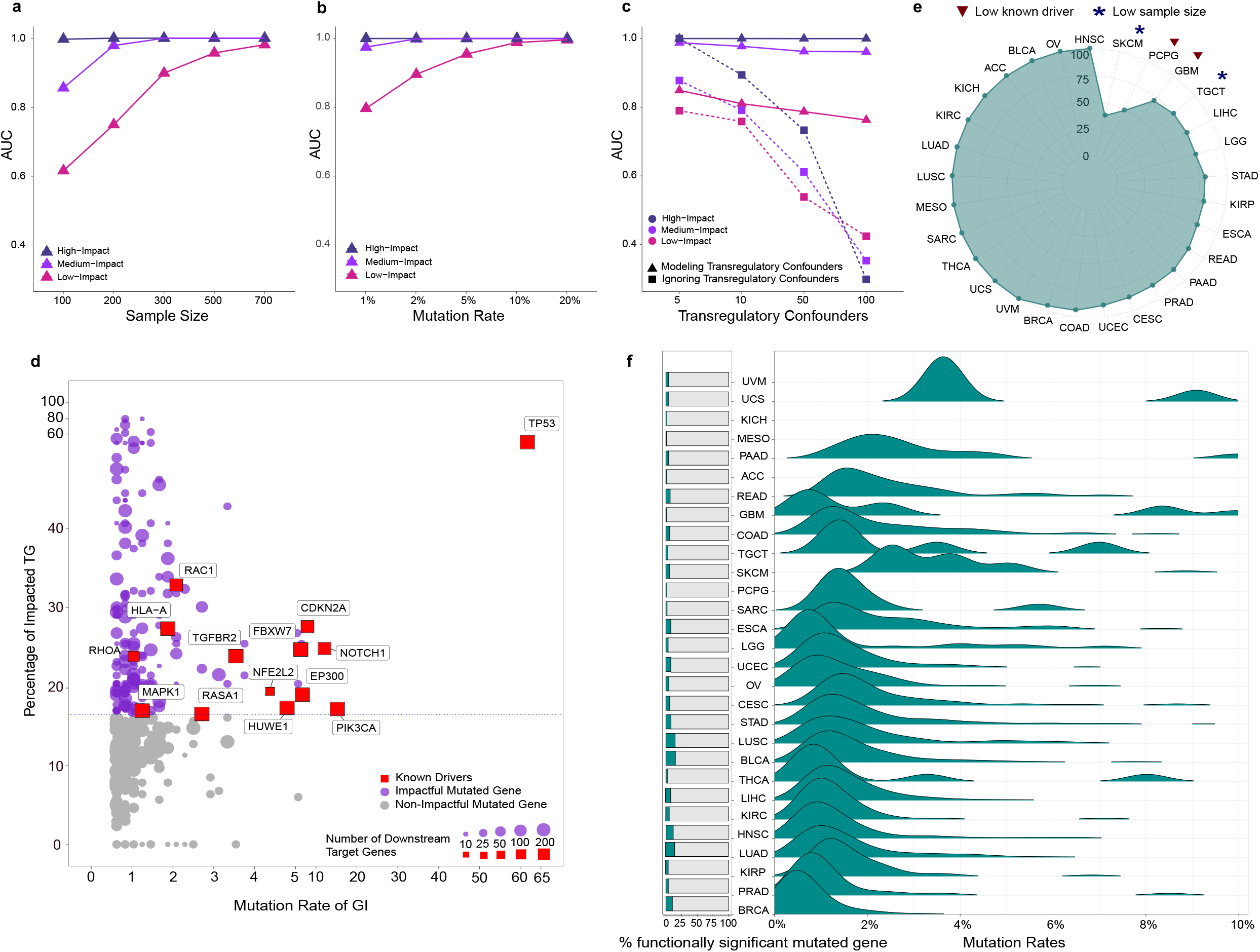
Computational benchmarking of SYS-Mut using simulation and PanCancer datasets. **a-c** Simulation-based benchmarking of the performance of SYS-Mut in detecting low, medium and high mutational impacts of the gene of interest across conditions including, (**a**) Number of samples available for training the model; (**b**) Mutation rates; and (**c**) Number of Trans-regulatory confounders. Note the significant performance degradation in mutational impact detection when the *trans*-regulatory confounders are not considered in the model (dashed-line). **d** Example of SYS-Mut based quantification of the functional impact of mutated genes in Head and Neck Squamous Cell Carcinoma (HNSC). The scatter plot denotes the fraction of impacted target genes associated with each indicated *GI* (Y-Axis), plotted against the *GI*’s mutation rate (X-Axis). The horizontal dashed line denotes the threshold of statistical significance (*P* < 0.05). Shown in purple are mutated genes with significant functional impact identified by SYS-Mut, with red nodes denoting previously-identified driver genes in HNSC. **e** The spider-plot indicates the percentage of previously-identified driver genes within each of 29 cancer types whose impact was recapitulated by SYS-Mut. Cancers where SYS-Mut detected impact in <75% of the previously-identified driver genes were due to either low sample size (*) or very low number of driver genes (∇). **f** Bar-graphs (Left) denote the percentage of mutated genes with significant transcriptional impact identified by SYS-Mut across cancers. Also shown in right, are the distributions of mutation rates of genes whose mutations were identified by SYS-Mut as exhibiting significant downstream impact.

### Application of SYS-Mut to interrogate mutational significance in a PanCancer cohort of solid tumors

Given the success of the above simulation studies, we next assessed for SYS-Mut’s ability to detect the downstream transcriptional impact of mutations in the challenging setting of multi-omics molecular profiling data derived from admixed primary tumor tissues. Accordingly, we applied SYS-Mut to analyze multi-omics molecular profiles obtained from the PanCancer genome atlas project [21], that included a total of ~8,000 tumor samples spanning 29 distinct cancer types (**Supplementary Table S2**). We first selected high-confidence somatic mutations per tumor sample identified using paired-analysis of tumor and matched-normal whole-exome sequencing data (see **Supplementary Information**). We then employed SYS-Mut to integrate somatic mutation calls with sCNA, DNAMeth and RNAseq profiles per tumor sample, thus estimating the impact of a *GI* to its downstream *TGs* (as an example, **Supplementary Table S3**), and the fraction of target genes whose expression levels were significantly impacted by mutations in their respective *GIs* within each of 29 cancer types (**Supplementary Fig. 4a-c**). As a control, we performed SYS-Mut based impact assessments of mutations in each GI on a set of random TGs with similar numbers of *trans*-regulatory confounders (**Supplementary Fig. 4d-e**), thus generating a *null distribution* of the fraction of TGs impacted by mutations in a given *GI*. Mutations in *GIs* that resulted in significantly (*P* < 0.05) higher fractions of impacted TGs as compared to the *null distribution* were deemed to be functionally significant (**Supplementary Fig. 4f**), thus resulting in the identification of functionally significant mutated *GIs* in each of the 29 distinct cancer types within the PanCancer dataset (**Fig. 2d and Supplementary Fig. 5**). Strikingly, SYS-Mut was able to successfully classify as functionally significant nearly all of the predicted cancer driver genes previously identified by integrating 26 distinct computational approaches applied to the PanCancer cohort (**Fig. 2e**) [19].

Indeed, SYS-Mut’s performance far exceeded that of Xseq (**Supplementary Table S4;** See **Supplementary Information**), a finding consistent with our simulation studies (**Fig. 2c**) that suggest the critical importance of modeling the transcriptional influence of cis-/trans-regulatory factors on the downstream transcriptional targets of mutated genes. The few instances where SYS-Mut did not detect significant functional impact of a previously predicted driver gene were either due to insufficient sample size or the lack of sufficient information regarding downstream transcriptional targets based on the current transcriptional influence network (**Fig. 2e**). In addition to being highly sensitive in detecting previously-identified cancer driver genes (**Fig. 2e**), SYS-Mut also identified rarely mutated genes with significant functional impact while maintaining a high degree of specificity by classifying on average less than 7% of all mutated genes as functionally significant across cancer types (**Fig. 2f**).

We next evaluated whether the mutated genes (*GIs*) that were deemed by SYS-Mut as functionally clustered within specific molecular sub-networks. We clustered *GIs* based on overlaps in their target gene identities, with *GIs* sharing 70~100% of their target genes in the transcriptional influence network being clustered together (**Supplementary Table S5**; See **Supplementary Information**). The resulting clusters were then ranked based on the number of cancer types in which their respective *GI*s were deemed by SYS-Mut to harbor functionally significant mutations (**Fig. 3a**). As expected, the top ranked clusters included *GI*s related to pathways and cellular processes known to be deregulated in cancer, including Cell Cycle, DNA Repair, TP53, TGF-β, and NF-κB (**Fig. 3a**).

**Fig. 3.**
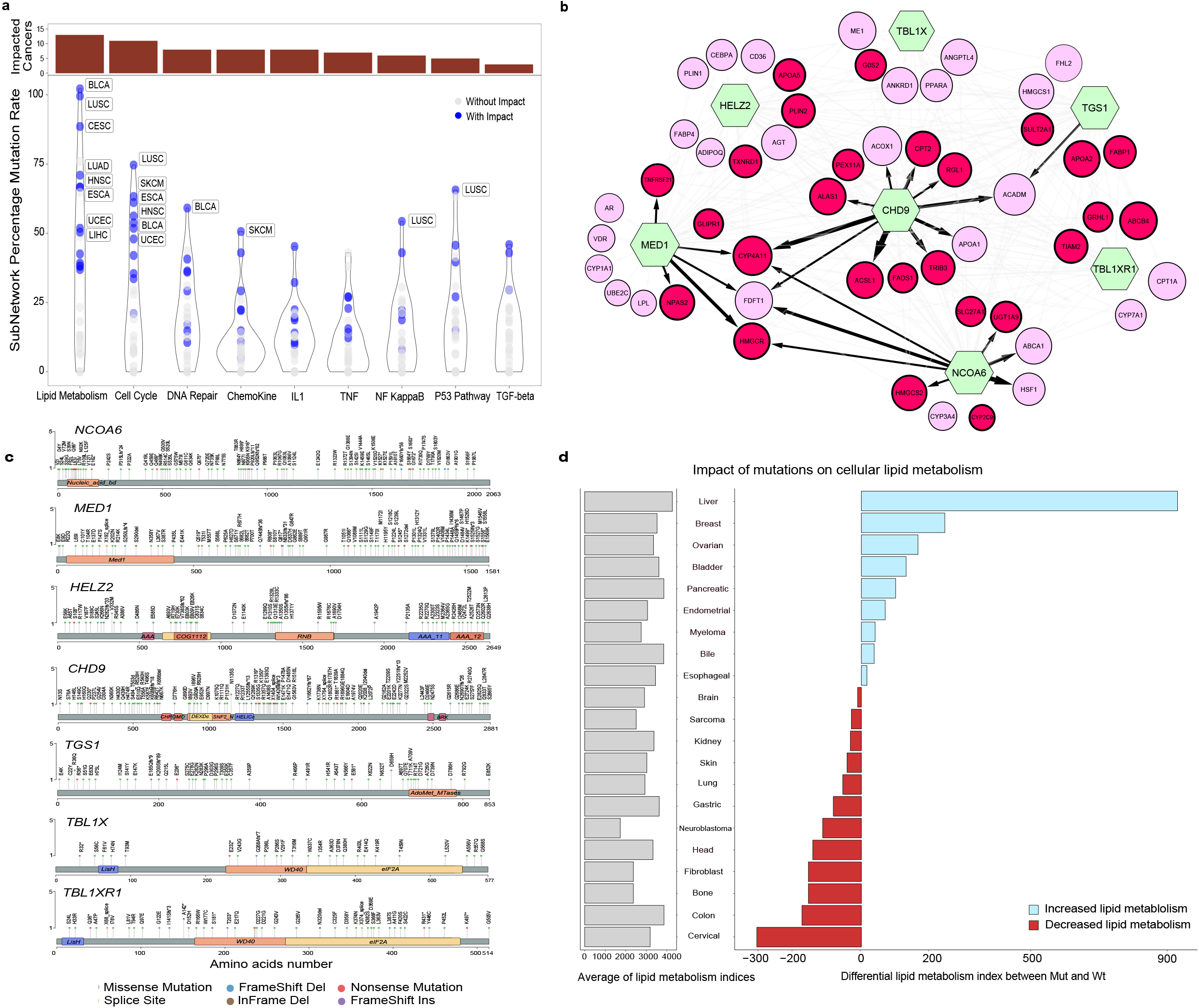
SYS-Mut based discovery of gene subnetworks harboring mutations with significant functional impact across cancers. **a** Violin plots denote the joint-mutation rate (Y-Axis) of sub-networks of genes identified by SYS-Mut as harboring mutations with significant downstream transcriptional impact across cancer types. Bar-graphs in dark red (top) denote the number of cancer types where SYS-Mut identified mutations in the subnetwork as exhibiting significant impact on downstream transcriptional programs. Note that SYS-Mut identified the lipid metabolism subnetwork as exhibiting significant mutational impact across the most number of cancer types. **b** Lipid metabolism gene sub-network detailing the convergent transcriptional impact of mutations in *GI*s (green hexagons) across cancer types in the PanCancer dataset. The size of the *TG* nodes (circles) corresponds to the number of cancer types where the *TG* was deemed as being impacted by mutation in the *GI*s. Red *TG* nodes (circles) are transcriptional targets uniquely impacted by *GI*s within this subnetwork, whereas pink *TG* nodes are also impacted by other previously-identified cancer driver genes. Directionality of the connection is indicated by arrows and the thickness of the arrows indicates the number of cancers where SYS-Mut identified a mutational impact of *GI* on the *TG*. The color of the arrow corresponds to the number of cancer types where SYS-Mut detected an impact of mutations in *GI* on the corresponding *TG* (black, corresponds to ≥4 cancer types, whereas light gray corresponds to <4 cancer types). **c** Lolliplot showing somatic mutations in the PanCancer dataset across the length of the amino-acid sequences for each of the *GI*s in the lipid metabolism network. Note the broad distribution of mutations throughout the length of the amino-acid sequences for each of the *GI*s in this network, a pattern suggestive of loss-of function as opposed to activating mutations. **d** Horizontal bar-plots indicating the impact of mutations on a transcriptional readout of lipid metabolism in the Cancer Cell Line Encyclopedia. Gray bars (left) indicate average lipid metabolism index across all cell lines in the cancer-type. Bars on the right detail increased (blue) or decreased (red) lipid metabolism in cell lines harboring mutations in the lipid metabolism network as compared to nonmutated cell lines.

Interestingly, the cluster that exhibited significant impact across the most number of distinct cancer types (**Fig. 3a**) included *TGS1, NCOA6, CHD9, MED1, HELZ2, TBL1X*, and *TBL1X1*. Strikingly, >89% of the *GIs* in this cluster and their associated *TGs* belong to the lipid metabolism pathway [22] (**Supplementary Table S6**). SYS-Mut also identified multiple *TG*s to be significantly impacted by more than one mutated *GI* within the lipid metabolism cluster (**Fig. 3b**), thus revealing how diverse mutational events can converge on a single cellular process. Importantly, none of these genes were identified as functionally significant by the pan-software [19] or Xseq frameworks (**Supplementary Table S4**). Next, in order to evaluate whether the non-synonymous somatic mutations in each of the mutated genes within the lipid metabolism cluster were likely to be activating or inactivating events, we assessed for potential mutational hotspots along the length of the respective proteins within the PanCancer TCGA dataset. However, we found somatic mutations to be broadly distributed along the length of each of the proteins within the lipid metabolism network with no evidence of mutational hotspots (**Fig. 3c**), thus suggesting that somatic mutations in these genes are more likely to be associated with loss-of-function as opposed to activating events. Consistent with this, a predefined gene expression based signature of lipid metabolic activity [22] revealed that cancer cell lines in 13 of 21 distinct tumor types represented in the CCLE dataset [23] were more likely to exhibit decreased lipid metabolic activity when they harbored mutations in the lipid metabolism network (**Fig. 3d;** See **Supplementary Information**).

### Mutations in lipid metabolism network are associated with poorer outcomes in Head and Neck Squamous Cell Carcinoma

Given that SYS-Mut identified non-synonymous somatic mutations in genes belonging to the lipid metabolism pathway as significantly impacting their downstream transcriptional targets across 13 cancer types of the PanCancer TCGA dataset (**Fig. 3a**), we proceeded to assess whether mutations in this pathway were associated with differential clinical outcomes in these specific cancer types (**Supplementary Fig. 6**). We observed a significant association between mutations in the lipid metabolism network and poorer Disease Specific Survival (DSS) only in Head and Neck Squamous Cell Carcinoma (HNSC) but not in any of the other cancer types (*P* = 0.028; **Fig. 4a**, **Supplementary Fig. 6**). Given that HPV-positivity is a known prognostic factor in Head and Neck Squamous Cell Carcinoma (HNSC) [24,25], we performed a subset analysis and found mutations in the lipid metabolism network to be significantly associated with poorer DSS even in HPV-Negative HNSC (*P* = 0.0036, **Fig. 4b**). Furthermore, the mutation rates of previously-identified HNSC driver genes [19] were similar between HNSCs mutant or wild-type for the lipid metabolism network (**Supplementary Fig. 7**). Strikingly, mutations in the lipid metabolism network were associated with poorer DSS in HPV-negative HNSC even after adjusting for clinical-pathologic variables such as stage and age (*P* = 0.0069, **Fig. 4b**), as well as mutations in known HNSC driver genes (*P* = 0.0123) [19]. These findings, taken together with SYS-Mut’s assessment of significant impact of mutations in the lipid metabolism network on downstream transcriptional targets (**Fig. 4b**), strongly suggest that genomic alterations targeting the lipid metabolism pathway are a functionally-significant molecular mechanism in HNSC.

**Fig. 4.**
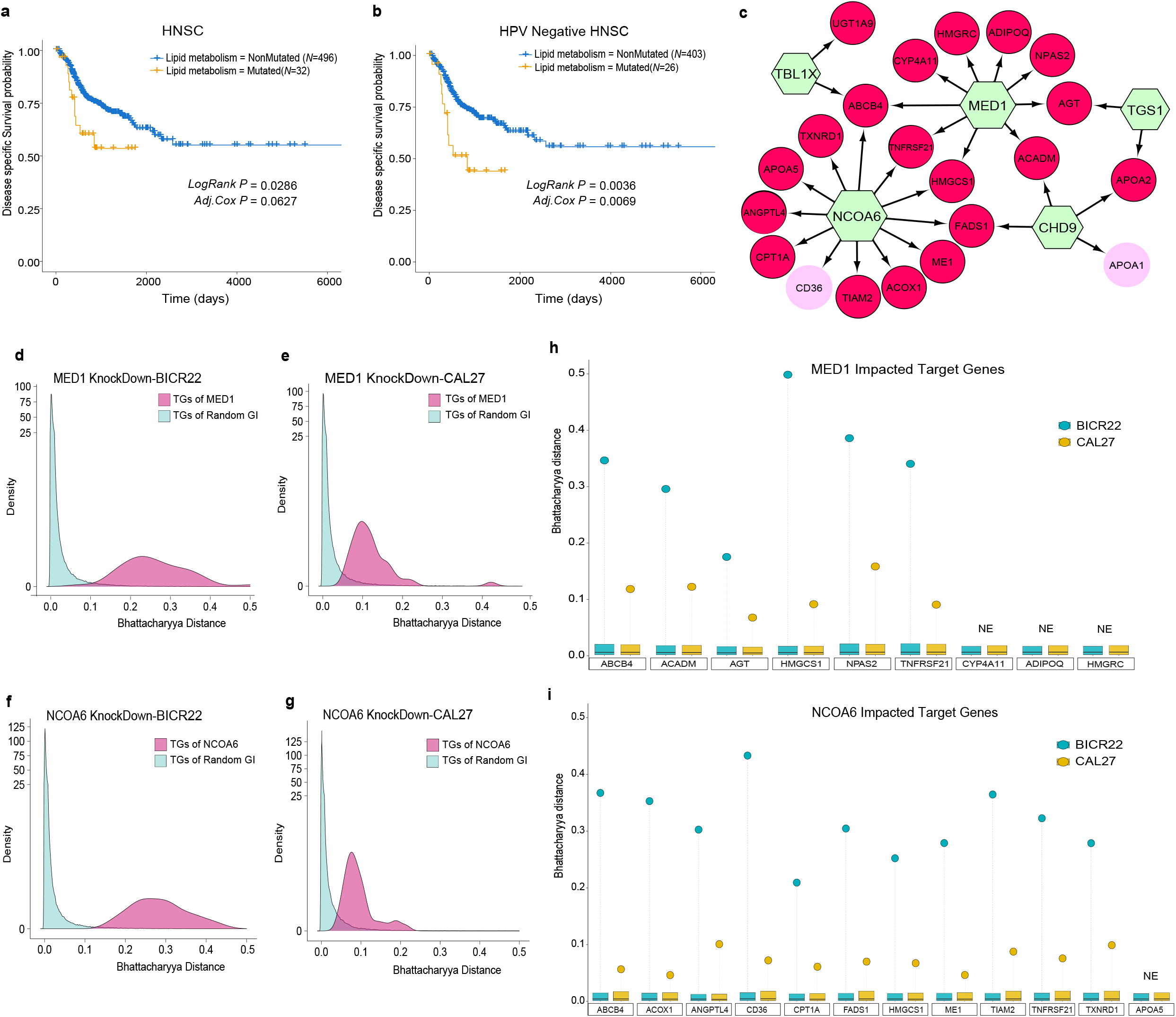
Impact of mutations in the lipid metabolism network in head and neck squamous cell carcinoma (HNSC). **a-b** Disease specific survival (DSS) in HNSC patients (**a**) or HPV-negative HNSC patients (**b**). Shown are Kaplan-Meier estimates of DSS for patients with tumors where one or more *GI*s in the lipid metabolism sub-network are Mutated (orange), as compared to their NonMutated (blue) counterparts. The statistical significance of differences in survival rates between Mutated and NonMutated categories was determined using the LogRank test (*LogRank P*); Also shown is the statistical significance estimated using the Wald test in a multivariable Cox regression model incorporating cancer stage and age (*Adj. Cox P*). **c** Lipid metabolism network detailing the convergent transcriptional impact of mutated *GIs* (green hexagon) in HNSC as identified by SYS-Mut. Red target gene (*TG*) nodes are transcriptional targets uniquely impacted by *GI*s in this subnetwork, whereas pink *TG* nodes are also impacted by other previously-identified driver genes in HNSC. **d-g** SYS-Mut based evaluation of the transcriptional impact of siRNA-based knockdown of *MED1* and *NCOA6* on their respective downstream target genes in distinct HNSC cell lines (BICR22 and CAL26). Shown are the distributions (magenta) of Bhattacharyya distances representing SYS-Mut’s assessments of the differential regulation of the downstream transcriptional targets of *MED1/NCOA6* in HNSC cell lines treated with either siRNA targeting *MED1* (siMED1) or *NCOA6* (siNCOA6) as compared to non-targeting control (siSCRAM). Also shown, as an internal control, are the distributions of Bhattacharyya distances (light blue) representing SYS-Mut’s assessment of impact of random untargeted *GI*s on their respective *TG*s. **h** and **i** depict the impact of *MED1* and *NCOA6* knockdown on target genes previously identified by SYS-Mut as being impacted by *MED1/NCOA6* mutations in HNSC primary tumors (as shown in panel **c**). Individual stems denote the Bhattacharyya distance (Y-Axis) corresponding to SYS-Mut’s assessment of the differential regulation of individual target genes (X-axis) upon *MED1/NCOA6* knockdown in BICR22 (dark cyan) and CAL27 (yellow) HNSC cell lines. The boxplots depict the distribution of Bhattacharya distances of randomly selected *GI*s on their respective *TGs*, thus serving as an internal control within each cell line model. *TG*s that are not expressed in a given cell line are denoted as NE.

The top two frequently mutated lipid metabolism associated genes (*MED1* and *NCOA6*) in HNSC did not exhibit any evidence of mutational hotspots (**Supplementary Fig. 8**), similar to our findings in the PanCancer context (**Fig. 3c**), thus suggesting that non-synonymous mutations in these genes are were more likely to be inactivating events. We therefore performed siRNA-based knockdown of *MED1* and *NCOA6* in two distinct HNSC cell lines (BICR22 and CAL27) that do not harbor somatic mutations in the lipid metabolism network, to functionally test the impact of inactivation of these genes on downstream transcriptional targets. Accordingly, we employed SYS-Mut to analyze the knockdown (siMED1 or siNCOA6) versus control (siSCRAM) RNASeq profiles in each cell line, followed by assessment of transcriptional impact on the transcriptional targets of the respective genes (See **Supplementary Information**). SYS-Mut revealed significant alterations in the regulatory influence of *MED1* or *NCOA6* on their respective target genes (*TG*s) upon siRNA-based knockdown, as evidenced by significantly higher Bhattacharyya distances of their respective *TG*s as compared to the Bhattacharyya distances of *TG*s of random untargeted genes in both BICR22 and CAL27 cell line models (**Fig. 4d-g**). SYS-Mut was indeed able to successfully recapitulate the previously-detected mutational impacts of *MED1* or *NCOA6* on their respective TGs (**Fig. 4c**) using the siRNA-based assessments in BICR22 or CAL27 cell lines (**Fig. 4h-i**). These functional assessments in distinct HNSC cell line models provide a strong support of SYS-Mut’s robustness in detecting the downstream transcriptional impact of mutations in genes even in molecular profiles derived from admixed primary tumor tissue samples.

### HNSC cells harboring mutations in lipid metabolism network exhibit differential sensitivity to glucocorticoid receptor agonists

Given that SYS-Mut identified mutations in the lipid metabolism network as impacting downstream transcriptional programs and survival in HNSC, we evaluated whether mutations in this network were associated with decreased lipid metabolism in head and neck cancer cell lines. Indeed, we found that mutations in genes belonging to the lipid metabolism network in head and neck cancer cell lines were associated with significantly lower lipid metabolic activity (**Fig. 5a**), a finding consistent with our position-based mutation assessments in primary tissues that also supported a likely inactivating function of mutations in these genes (**Supplementary Fig. 8**).

**Fig. 5.**
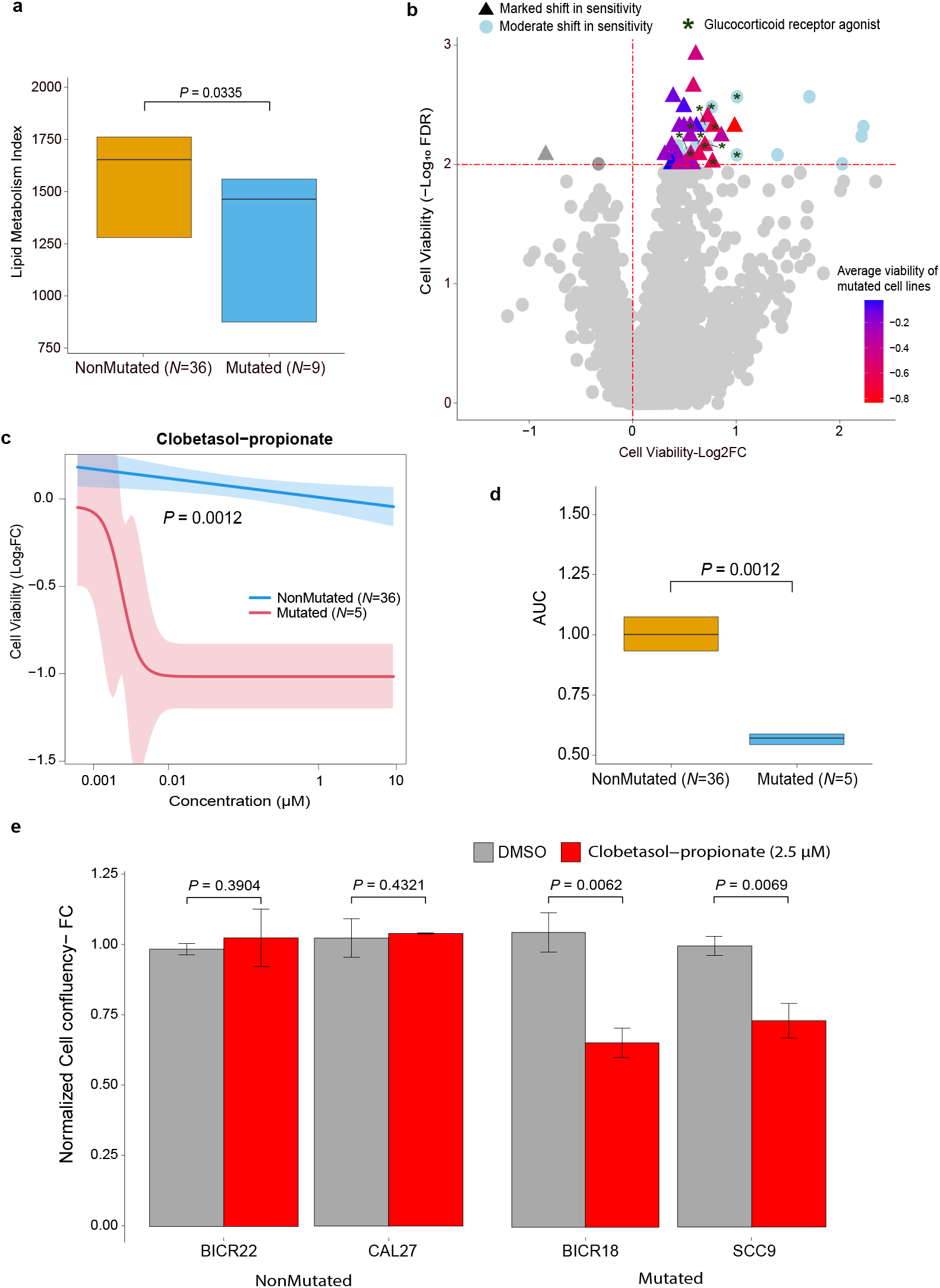
Differential drug sensitivity of HNSC cell lines harboring mutations in the lipid metabolism network as compared to their nonmutated counterparts. **a** Boxplots detailing lipid metabolic activity in HNSC cell lines harboring mutations in the lipid metabolism network (Mutated) as compared to their NonMutated counterparts. Statistical significance of differences in lipid metabolic activity was estimated using a Wilcoxon test. **b** Volcano plot depicting significant differential sensitivity between Mutated and NonMutated HNSC cell lines across drugs in the PRISM primary screen. X-axis denotes the difference (log_2_fold-change) in average cell viability between Mutated and NonMutated HNSC cell lines upon drug treatment, whereas the Y-Axis denotes the statistical significance (-log_10_FDR) of the difference assessed using a Wilcoxon test. Points above the horizontal red dashed line denote drugs inducing a significant shift (log_10_<0.01) in average cell viability between Mutated and NonMutated HNSC cell lines. Drugs were additionally denoted as exhibiting Marked (triangles) shift in sensitivity if the Mutated cells exhibited, on average, lower cell viability upon drug treatment as compared to DMSO control (indicated by negative log_2_ cell viability in the PRISM primary screen) while the NonMutated cells were non-responsive (indicated by positive log2 cell viability values in the PRISM primary screen). Glucocorticoid receptor agonists are highlighted with dark green stars. **c** Average clobetasol-proprionate dose-response curves for Mutated and NonMutated (blue) HNSC cell lines derived from the PRISM secondary screen dataset. Y-axis denotes the difference in cell viability (Log_2_fold change) upon treatment with different doses (X-axis) of clobetasol-proprionate as compared to DMSO control. **d**. Boxplots detailing the comparison of Area Under the Curve (AUC) estimates for clobetasol-propionate dose-response curves in Mutated versus NonMutated HNSC cell lines obtained from the PRISM secondary screen dataset. Statistical significance was assessed using a Wilcoxon test. **e** IncuCyte-based cell growth assessments in Mutated (SCC9, BICR18) and NonMutated (BICR22, CAL27) HNSC cells treated with clobetasol-propionate (2.5 μM) as compared to DMSO control treatment. Y-axis represents the average fold-change in cell confluence values at the final timepoint (120hrs for SCC9; 96hrs for BICR22, CAL27 and BICR18) normalized to DMSO control, plotted as mean ± SD, obtained from at least 3 replicate experiments. The statistical significance of differences in normalized cell confluency between the respective test versus control groups were estimated using a Student’s t-test assuming unequal variances.

We next proceeded to orthogonally assess the functional significance of mutations in these genes by analyzing drug efficacy data published as part of the PRISM drug repurposing resource within the DepMap portal [26] (See **Supplementary Information**). Specifically, we ranked drugs in the PRISM primary screen based on whether HNSC cells harboring mutations in lipid metabolism exhibited significant differential sensitivity to the drug as compared to non-mutated HNSC counterparts. Interestingly, the top ranked drugs within the PRISM primary screen that exhibited differential sensitivity between mutated versus non-mutated HNSC cell lines were significantly enriched for glucocorticoid receptor agonists (*P* < 1.5×10^-13^; **Fig. 5b** and **Supplementary Table S7**). Indeed, HNSC cell lines harboring mutations in the lipid metabolism network were more likely to exhibit markedly higher sensitivity to glucocorticoid receptor agonists as compared to their non-mutated counterparts (**Fig. 5b**). Given that the primary PRISM screen employed a single drug concentration (2.5□μM) across all drugs, we analyzed the PRISM secondary screen dataset within DepMap that assesses cell viability for a subset of drugs (*N* = 1448) and cell lines across doses ranging from 0.0006-10μM. Here too, we found clobetasol-propionate, a glucocorticoid receptor agonist, as exhibiting the strongest differential sensitivity between mutated and non-mutated HNSC cell lines (*P* = 0.0013; **Fig. 5c, 5d** and **Supplementary Table S8**). Given the known role of the glucocorticoid receptor in regulating cellular lipid metabolism [27–29], we performed an in-house validation study to test whether clobetasol-propionate differentially impacts the growth (See **Supplementary Information**) of distinct HNSC cell lines that are either mutated (BICR18, SCC9) or non-mutated (BICR22, CAL27). Of note, neither of the mutated cell lines were included in the secondary PRISM screens, thus enabling us to additionally test for the generality of our initial findings. Strikingly, and consistent with our findings in the PRISM drug screen, we observed that the mutated HNSC cell lines exhibited significant reduction in cell growth as compared to DMSO control, while the non-mutated cell lines were unresponsive to the drug (**Fig. 5e**). Taken together, these findings strongly support SYS-Mut’s assessment that mutations in the lipid metabolism network exhibit significant functional impact in Head and Neck Cancer (**Fig. 3, 4**).

## Discussion

Individual tumors typically harbor multiple genomic aberrations in addition to alterations spanning the epigenomic and transcriptomic scales, thus posing a significant challenge to distinguish functionally significant mutations from the background of passenger events [4]. In addressing this challenge, we developed a multi-omics integrative systems biology framework, SYS-Mut, to decode the functional significance of somatic mutations in admixed cancer tissue samples. SYS-Mut is designed to quantify the functional impact of mutations on downstream transcriptional programs using a rigorous Bayesian regression framework that accounts for both *cis-* and *trans*-regulatory genomic, transcriptomic and epigenetic factors (**Fig. 1; Supplementary Table S3**).

Computational benchmarking using simulated data demonstrated the robustness of SYS-Mut in decoding the impact of mutations on downstream transcriptional deregulation even in the presence of co-occurring transcriptional and genomic aberrations similar to real-world tumor molecular profiles (**Fig. 2**). This key capability is foundational to SYS-Mut’s ability to robustly detect mutational impact even in the context of rare events confounded by a large number of cis-/trans-regulatory factors (**Fig. 2c, 2d**), thus distinguishing SYS-Mut from previous approaches such as Xseq [15]. Indeed, our application of SYS-Mut to analyze the impact of mutations in a PanCancer multi-omics dataset [20] recapitulated the vast majority of previously-identified cancer driver genes [19] across cancer types, while additionally enabling prioritization of a relatively small fraction (<7%) of rarely mutated genes that exhibit significant transcriptional impact within each cancer-type (**Fig. 2e**). Importantly, SYS-Mut models the transcriptional impact of mutations by accounting for co-occurring molecular aberrations within each tumor sample, thus uniquely enabling it to decode the tissue-specific functional significance of rare mutational events (**Fig. 3a**). SYS-Mut identified recurrent mutations in the lipid metabolism subnetwork to be functionally significant only in a subset of cancer types (**Fig. 3a**), thus highlighting its ability to contextualize the impact of mutations within the tissue-specific molecular milieu.

In addition to determining the functional significance of a rarely mutated genes, SYS-Mut also provides insights into the key downstream transcriptional targets that are impacted by mutations in the gene. Indeed, SYS-Mut’s specific predictions of downstream transcriptional impact assessments were orthogonally validated using genetic manipulation studies in cancer cell line models (**Fig. 4d-4i** and **Supplementary Fig. S9**), thus further supporting the robustness and utility of this systems biology integrative framework.

The translational utility of SYS-Mut is evidenced by its assessments in the PanCancer multi-omics dataset that revealed rare mutations in a lipidogenic subnetwork as being functionally significant in head and neck cancer (**Figs. 3 & 4**). Our findings of somatic mutations in the lipid metabolism subnetwork that result in lower lipid metabolic activity and poorer outcomes in HNSC (**Figs. 3-5**) are consistent with a prior report detailing lower lipid metabolism as being associated with poorer survival outcomes[22]. These findings are further supported by our findings that head and neck cancer cells harboring mutations in the lipid metabolism network are more likely to exhibit reduced lipid metabolic activity and are sensitive to glucocorticoid receptor agonists (**Fig. 5b-e**). While glucocorticoid receptor signaling is known to play a tumor-suppressive role in a subset of cancers [30–32], this is the first study to show that glucocorticoid receptor agonists can suppress the growth of head and neck cancer cells harboring mutations in the lipid metabolism pathway. While the precise molecular mechanism underlying this differential sensitivity to glucocorticoid receptor agonists remains to be discovered, further investigation is warranted to determine whether targeting this signaling axis might benefit a subset of HNSC patients.

A unique advantage of SYS-Mut is that the hierarchical Bayesian regression model is naturally scalable from the current mutated versus nonmutated assessments to quantify the differential impact on target gene expression profiles across multiple user-defined subgroups of tumor samples. As such, SYS-Mut can be easily extended to assess for the differential impact of multiple categories of mutations in a given gene, such as mutations in different functional domains of a given protein. Conversely, SYS-Mut can also be employed to assess for functional synergies by defining subgroups of tumors characterized by combinations of mutational events across genes. While SYS-Mut is only able to assess the functional impact of mutations in genes represented within the transcriptional influence network (**Supplementary Table. S1**), the completeness and accuracy of the underlying signaling networks will continue to improve, thus expanding SYS-Mut’s capabilities. Nonetheless, we anticipate future versions of SYS-Mut to also incorporate phosphoproteomic measurements thus enabling assessments of the impact of mutations on downstream signaling networks.

## Conclusions

In conclusion, we propose SYS-Mut as a robust, flexible and scalable systems biology approach that can reveal the functional significance of the growing number of rare mutational events identified by large-scale cancer sequencing initiatives. Our study highlights the utility of combining regulatory network connectivity with multi-omics data to unravel rare but functionally significant mutations in cancer thus allowing for the discovery of novel therapeutic targets in this complex disease.

## Supplementary Information

### SYS-Mut Model

In order to account for potential nonlinear relationships between the *cis-*regulatory measurements and gene expression levels, SYS-Mut employs a centered sigmoid and soft thresholding function, to transform the DNA Methylation and sCNA data, respectively.

### Hierarchical Bayesian inference details

Accordingly, SYS-Mut employs a Bayesian inference framework that learns the relationship between observed data and all relevant influencing parameters. The data are considered to be a sample randomly drawn from the sampling distribution p(***y*** | ***x***, **Θ, Ψ**). Bayesian model estimates full posterior distributions of the model parameters, and it incorporates a prior knowledge about the model parameters using hierarchical Bayesian models. The likelihood function *l*(**Θ, Ψ**; ***y***) required for the application of Bayes theorem equals the following sampling distribution p(***y*** | ***x***, **Θ, Ψ**). Notably, a hierarchical prior is used instead of selecting a prior from a single family of distributions (**Supplementary Fig. 3a**).

As such, SYS-Mut constructs the following model, where the subscripts *TG*, *GI* and *PG* are omitted for notation convenience:

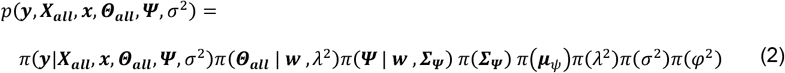

where

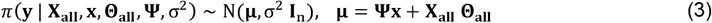

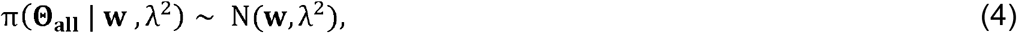

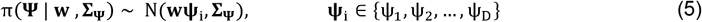

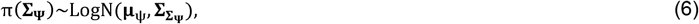

Where **∑_Ψ_** denotes the cluster covariance matrix.

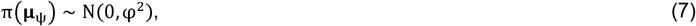

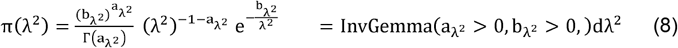

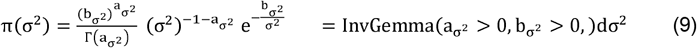

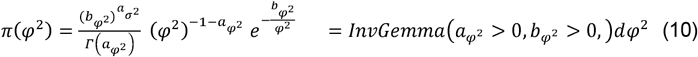

Gibbs sampling, as implemented in JAGS[33], a program for Bayesian inference using Markov chain Monte Carlo (MCMC), is applied to obtain the approximate posterior distributions for the model parameters **Ψ, Θ_all_**, thus providing the basis to assess for the functional effects of mutations in GI on TG.

### Building a transcriptional influence network

SYS-Mut builds a genome-scale transcriptional influence network capturing the regulatory roadmap of gene transcription by integrating signaling and transcription factor network information from the several publicly available datasets including TRED, TRRUST, and Paradigm Super-Pathway. TRED, a transcriptional regulatory element database and a platform for *in silico* gene regulation studies, which is an integrated repository for both *cis*- / *trans*-regulatory elements in mammals (available at http://rulai.cshl.edu/TRED) [34]. TRRUST, transcriptional regulatory relationships unraveled by sentence-based text mining dataset, which is a manually curated database of human and mouse transcriptional regulatory networks (available at https://www.grnpedia.org/trrust/) [35]. Paradigm Super-Pathway network, a set of interactions from several sources including NCI-PID, Reactome, and KEGG, and superimposed them into a single network (Super-Pathway). The Super-Pathway contained 7,369 proteins, 9,354 multi-protein complexes, 2,092 families, and 592 cellular processes connected by 45,315 interactions (available at https://github.com/epaull/TieDIE/tree/master/pathways) [14].

The 18,651 unique nodes of the network consisting of *Gene*, *Complex formation, Abstract, Family* are connected via 120655 interactions of *Transcriptional regulation* (-*t*), *Post-transcriptional activation* (-*a*), *Member, and Component*. In addition to the built transcriptional influence network, any such influence network encoding gene regulation can be used as the input of the SYS-Mut.

### Simulation comparison

We have generated a large set of synthetic data for mutation, gene expression, sCNA and DNAMeth. We constrained the input data to mimic the distributions of normalized multi-omics data within TCGA PanCancer dataset [20]. A variety of noise levels sampled from a normal distribution 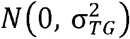, were added to the data to mimic measurement noise as well as unknown extrinsic influences on gene expression. A variety of conditions including, number of samples (100, 300, 500, and 700) available for training the model, different mutation rates of *GI* (0.5%, 1%, 2%, and 5%), and number of *trans*-regulators of *TG* (5, 10, 50, 100) that could confound the impact mutation assessment were created for the comparison. Additionally, a parameter of mutational impact was applied to the mutated samples, aiming to simulate three levels of low, medium, and high impacts. We then applied SYS-Mut model to the synthetic data for 100 iterations, and the posterior parameters of each simulation were estimated. The level of impact was estimated by statistical comparison of two posterior distributions of the Mutated versus Nonmutated samples. In particular, the significance impact level of a *GI* is represented by a t-test *P*-value of comparison between the two distributions.

### Preprocessing and normalizing the PanCancer multi-omics data streams

We conducted comprehensive integrative molecular analyses on ~8000 samples spanning 29 solid tumor types of the cancer genome atlas (TCGA) dataset [21]. In particular, we collected multi-omics data consisting of DNA methylation, somatic copy number alterations, RNA sequencing gene expression, somatic mutation calls, and clinical data from the PanCancer atlas genome portal (https://gdc.cancer.gov/about-data/publications/PanCan-CellOfOrigin) [21].

Specifically, Infinium human DNA methylation profiles were obtained from a combined and further probe-by-probe normalized data from two generations of Infinium arrays, HumanMethylation27 (HM27) and HumanMethylation450 (HM450), including 22601 probes shared between two platforms. The level 2 DNA methylation, aligned to the GRCh38(V.28) reference genome[36], were preprocessed using our pipeline and the average of all mapped probes associated to a gene were considered as DNA methylation value of the gene. For some probes, which associated with more than one gene, we assigned the beta values to all the corresponding gene symbols. Moreover, we processed somatic copy number alterations (sCNA) data, assessed as deviations in the tumor sample from the paired normal tissue sample. The sCNA profiles were generated based on Affymetrix SNP6 array copy number inference pipeline [37] and the preliminary copy number at each probe locus was inferred by Birdseed analysis of raw files [38]. The probs locus with the length <50bp were filtered in our analysis as they are considered as artifacts. To performed a gene level CNA analysis, we aligned the probes and their corresponding segment to the genomic locations provided in the GRCh38(V.28) reference genome [36]. We Also, employed a log_2_-transformed Fragments per kilo-base of mRNA per million reads (FPKM) of the estimated gene expression profiles (level 3, from “IlluminaHiSeq_RNASeqV2” platform) of tumor samples from the TCGA normalized read counts, which aligned to the GRCh38(V.28) reference genome [36].

### Identification of high confidence somatic mutations in the PanCancer dataset

Somatic single-nucleotide mutation calls (SNVs and indels), identified in whole-exome sequencing (WES) studies of the 29 solid tumor types, were obtained from PanCancer atlas platform (MC3 V.0.2.8, https://synapse.org/MC3) [21]. The MC3 effort provided consensus calls of variants from multiple genomic platforms including, MuTect, VarScan2, Somatic Sniper, MuSE, Radia, Indelocator, and Pindel. We extracted the position and nucleotide change information for all single-nucleotide somatic mutations, which integrated into a combined MAF file. We then categorized aberrations as synonymous (*Wt*) and non-synonymous (*Mut*). Missense, Frame shift deletion, Nonsense, Inframe deletion, Inframe insertion, Splice site, Frame shift insertion, Translation start site, Nonstop mutations were marked as non-synonymous (*Mut*) mutations, while Silent, 3′UTR, RNA, Intron, 3′Flank, 5′UTR, 5′Flank were labeled are synonymous (*Wt*) group. We additionally annotated a single unique mutation entry having multiple annotations belong to different variant classification as non-synonymous, where at least one of the multiple entries of the annotations shows in the non-synonymous category. We performed extensive filtering to minimize sequencing artifacts, mutation calling errors and to exclude likely false-positive mutations. In fact, mutation calls with tumor depth or normal depth less than 30 reads, and tumor frequency less than 0.1 or normal frequency greater than 0.05 were filtered out from our analysis. We also excluded hypermutated samples, as they likely reflect distinct underlying mutational processes, and they tend to have an adverse effect on statistical power. We identified hypermutated samples as having more mutations than 1.5 times the interquartile range above the third quartile of samples within the same cancer cohort. Moreover, in colorectal squamous adenocarcinoma (COAD), samples with high microsatellite instability (MSI-H) [39], were removed from the analysis as they have distinct clinicopathological characteristics consisting of factors predicting positive and negative outcomes.

### Quantitative analysis of SYS-Mut across 29 solid tumors of TCGA

A consensus list of cancer driver genes identified by integrating 26 distinct computational approaches applied to the PanCancer cohort [19] was used to benchmark SYS-Mut performance. Some of the previously identified driver genes were filtered out due to either the lake or deficiency of transcriptional connection in the influence network (<5 transcriptional connections from a *GI* to its *TGs*) or deficiency of mutated samples (<3 samples with somatic mutation in the *GI*) in a cancer tissue. The PanCancer RNAseq and sCNA data across 29 solid tumors were additionally anlayzed using Xseq (R Package V. 0.2.1). Xseq was run with default parameters on the PanCancer cohort after mapping the mutations in each of the tumor samples into the Xseq mutation classes. Xseq generated a likelihood of Impact, *P*(*D*) for each mutated gene in a specific cancer type while also inferring the likelihood of mutational impact per tumor sample, *P*(*F*).

### Integrative functional analysis of *GIs* identified by SYS-Mut as functionally significant

To perform integrated functional analysis across all tumor types, we evaluated whether the mutations identified by SYS-Mut as functionally significant, jointly defined a unique network of cancers. Given the very low mutation rates in many of the GIs identified by SYS-Mut as functionally significant, we defined clusters of GIs that shared %70~100 of their target genes in the transcriptional influence network, using ward2 Hierarchical clustering method, where it creates the groups such that variance is minimized within clusters. 316 unique clusters were created, while some of the clusters include only one *GI*.

### Analysis of the survival data across the solid tumors

The clinical data were downloaded from the data portal of PanCancer atlas genome project [21]. The TCGA samples barcode structure enables integration of patient-based clinical data with sample-based molecular data. Clinical outcome of disease-specific survival (DSS) across the solid tumors were analyzed for association of mutations in the 7-genes (*TGS1, NCOA6, CHD9, MED1, HELZ2, TBL1X*, and *TBL1X1*) of lipid metabolism network using Kaplan–Meier estimates. The statistical significance of differences in survival rates between Mutated and NonMutated categories was determined using the LogRank test. Additionally, Hazard ratios and the corresponding *P*-values were calculated from a Cox proportional hazards model.

### Assessment of lipid metabolic activity in HNSC cell lines

Gene expression profiles and the corresponding mutation annotation file of 40 head and neck cancer cell lines of the cancer cell line encyclopedia (CCLE) dataset were obtained from DepMap portal [26]. After removing the Hypermutated cell lines, we assessed lipid metabolic activity on the 32 remaining HNSC cell lines. A 7-gene of lipid metabolism signature [22] (*TGS1, NCOA6, CHD9, MED1, HELZ2, TBL1X*, and *TBL1X1*), was used as a surrogate estimate. For each of 32 HNSC cell lines, a lipid metabolism index was estimated using the single-sample gene set enrichment (ssGSEA) methodology [40]. Wilcoxon *P*-value of the comparison between two group of the cell lines harboring mutations in the 7-genes of lipid metabolism network (Mutated) as compared to their NonMutated counterparts were calculated.

### Cell culture

HNSC cell lines BICR18 (cat# 06051601) and BICR22 (cat#04072106) were purchased from Sigma Aldrich while CAL27 (cat# CRL-2095) and SCC9 (cat# CRL-1629) were purchased from American Type Culture Collection (ATCC, Manassas, VA). BICR 18 and BICR22 cell lines were cultured in Dulbecco’s Modified Eagle Medium (DMEM) (Gibco, cat# 11-875-119) supplemented with 2% fetal bovine serum (FBS) (Gibco, cat# A3160501) +0.4 micrograms/ml Hydrocortisone (Selleckchem, cat# S1696). CAL27 was cultured in Dulbecco’s Modified Eagle Medium (DMEM) supplemented with 10% fetal bovine serum. SCC9 was maintained in 1:1 Dulbecco’s modified Eagle’s medium and Ham’s F12 (Gibco, cat# 11-320-033) supplemented with 10% FBS +0.4 micrograms/ml Hydrocortisone. Cell lines were screened periodically for mycoplasma contamination using the MycoAlert kit (Lonza, Basel, Switzerland, # LT07-518).

### siRNA knockdown

HNSC cell lines were seeded in triplicate and transfected with 50nM siRNAs directed against MED1 (Dharmacon, cat# L-004126) and NCOA6 (Dharmacon, cat# L-019107) or non-targeting/control siRNA (Dharmacon, cat# D-001810), in serum-free media using RNAiMAX (Life Technologies, #13778030) transfection reagent. Cells were harvested after 48 hours and RNA isolated as per manufacturer’s protocol (Qaigen cat# 74134) and submitted for RNA-sequencing, resulting in an average of 60 million paired-end reads per replicate across cell line models and siRNA treatments.

We generated and annotated the RNASeq expression read counts by first aligning to the GRCh38.p13 reference genome using RNASeq Aligner[41] and conducting the transcriptome assembly using the Cuffquant from the Cufflinks assembler and abundance estimation algorithms[42]. The RNASeq expression read counts and normalized expression levels (FPKM) for each gene, transcript, TSS group, and CDS group in the experiment are generated using the Cuffnorm[42,43]. After removing the duplicated gene symbols by selecting the genes with the maximum mean absolute deviation we then normalized and transformed the expression levels of each gene in to log_2_ scale.

To identify the impact on the transcriptional down steam target gens, we employed SYS-Mut to analyze the siMED1-knockdown and siNCOA6-knockdown versus control (siSCRAM) RNASeq profiles within each cell line, independently. The Bhattacharyya distances of si***X***, where ***X*** = {*MED1, NCOA6*}, and siSCRAM (*D_ψsiX, ψsiSCRAM_*), were calculated within each cell line over the estimated posterior distributions for the expression of the transcriptional targets (*TGs*) of *MED1* and *NCOA6*. To identify the control, SYS-Mut then applied to all other *GI*s with similar number of confounders. The Bhattacharyya distances of si***Y*** and siSCRAM (*D_ψsiY, ψsiSCRAM_*), where ***Y*** = {all other genes with similar number of confounders}, were also calculated within each cell line, independently. SYS-Mut then assessed the regulatory influence of *MED1, NCOA6* on their *TGs* by asking whether *D_ψsiX, ψsiSCRAM_* where ***X*** = {*MED1, NCOA6*}, is significantly greater than distribution of all the control Bhattacharyya distances in both BICR22 and CAL27 cell line models.

Moreover, principal component analysis (PCA) of RNASeq-based whole-transcriptome profiles derived from two distinct HNSC cell lines (BICR22 and CAL27) after 48-hrs of treatment with either siRNA targeting *NCOA6* (siNCOA6) or *MED1* (siMED1), as compared to non-targeting control (siSCRAM) was performed. Note the biologic replicates from each cell line and treatment category clustering together.

### *In silico* drug sensitivity screen in HNSC cell lines

In order to evaluate potential novel therapeutic targets correlated with mutated lipid metabolism pathway, we assessed the drug efficacy over the identified rare driver-genes of lipid metabolism pathway. Chemical perturbation log_2_fold-change data, calculated relative to DMSO, from the PRISM repurposing primary screen of 4518 drug, performed at 2.5□μM, on the HNSC cell lines of DepMap portal[26] were analyzed. Lipid metabolism pathway were mutated in 5 out of 28 HNSC cell lines. A comparison of the Wilcoxon *P*-value, average cell viability log fold-change data, and fisher *P*-values for the cell lines that harboring mutations in lipid metabolism versus the wild-type cell lines were performed. We then analyzed and selected the significant primary PRISM repurposing drugs based on Wilcoxon test (*P* <0.01) of comparison between two group of the cell lines. The significant drugs with the same mechanism of actions (moa) are grouped and ranked based on marked and moderate shift in their cell viability and the corresponding fisher *P*-values.

Additionally, we employed the PRISM secondary screen chemical perturbation log fold-change data for the HNSC cell lines in DepMap that also have drug sensitivity data reported as areas under the curve (AUC) of the dose-response ranges of 0.0006-10μM. Similar to the primary screen, we have assessed the drug efficacy over the identified rare driver-genes of lipid metabolism pathway upon the two group of the HNSC cell lines. The average AUCs of the cell lines that harboring mutations in lipid metabolism versus the wild-type cell lines were calculated. We then analyzed and ranked the significant secondary PRISM repurposing drugs based on Wilcoxon *P*-value of comparison between two categorize of the cell lines.

### Cell proliferation assay

For the drug experiments, Cells were seeded at densities of 1000 cells/well for BICR18 and SCC9 or 5000 cells/well for BICR22 and CAL27 in a 96-well plate and maintained in culture overnight in respective culture media. 24 hours later, cells were treated with 2.5μM Clobetasol Propionate (cat# S2584), in culture media supplemented in 2% FBS, using DMSO as a control (Tocris, cat# 3176). Treated cells were then imaged on the IncuCyte ZOOM automated live cell kinetic imaging system (Essen Bioscience, Ann Arbor, MI) for a course of 4-5 days with drug-treated media replenished every 3 days and assessed for confluence as per the manufacturer’s protocol.

### Availability of data and materials

SYS-Mut has been implemented as a set of interoperable and modular R scripts that can be run on either a stand-alone server or on a high-performance computing cluster. The SYS-Mut package, related tutorials, as well as the generated or analyzed data underpinning the analyses in the paper can be downloaded from GitHub at https://github.com/skhalighicase/SYS-Mut. Other software packages and bioinformatics tools used in this study are indicated in the GitHub page. Data on intermediate results as well as the code to generate specific analyses are also available at the same address.

## Funding

This research was supported by PHS awards: T32 CA094186 (S. Khalighi), K25 DK115904 (V. Varadan), P30 CA043703 (V. Varadan), P20 CA233216 (V. Varadan). U54 CA163060 (K. Guda, V. Varadan), Case GI SPORE P50 CA150964 (K. Guda, V. Varadan), DeGregorio Family Foundation (K. Guda).

## Competing interests

V. V. is currently employed at Merck Research Laboratories. The authors declare no other competing interests.

## Notes

https://github.com/skhalighicase/SYS-Mut

### Competing Interest Statement

Vinay Varadan is currently employed at Merck Research Laboratories. The authors declare no other competing interests.

