## Supplemental Figures for "SYS-Mut: Decoding the Functional Significance of Rare Somatic Mutations in Cancer"

**Supplementary Figures**

**Supplementary Fig 1.** Fraction of known drivers amongst recurrently mutated genes across diverse cancer types.

**Supplementary Fig 2.** Schematic illustration of the regulatory interactions of a mutated gene (GI) and its downstream transcriptional target genes (TGs).

**Supplementary Fig 3.** Detailed methodology to identify mutated genes with significant impact in primary tumor datasets.

**Supplementary Fig 4.** Network based summarization of mutational impact of GI using a network randomization strategy.

**Supplementary Fig 5.** Quantitative analysis of 29 solid tumors of TCGA.

**Supplementary Fig 6.** Association of mutations in the lipid metabolism network across cancer types with disease specific survival.

**Supplementary Fig 7.** Non-exclusivity of mutations in lipid metabolism network and previously-known HNSC driver genes.

**Supplementary Fig 8.** Mutational hotspot assessments for MED1 and NCOA6 in Head and Neck Squamous Cell Carcinoma.

**Supplementary Fig 9.** RNAseq profiling of distinct HNSC cell lines upon siRNA-based knockdown of MED1 and NCOA6.

### Supplementary Tables

**Supplementary Table S1.** Transcriptional influence network employed by SYS-Mut.

**Supplementary Table S2.** Number of tumor samples analyzed by SYS-Mut within each cancer type in the pan-cancer multi-omics dataset.

**Supplementary Table S3.** Example of SYS-Mut estimated output for each GI to TG in head and neck cancer.

**Supplementary Table S4.** Comparison of SYSMut Output with Xseq method.

**Supplementary Table S5.** Clusters of mutated genes with significant functional impact across cancers.

**Supplementary Table S6.** SYS-Mut identified subnetwork with significant impact across cancers is predominantly populated with genes associated with lipid metabolism.

**Supplementary Table S7.** PRISM primary screen detailing the mechanisms of action of drugs exhibiting significant shifts in sensitivity between Mutated and NonMutated HNSC cell lines.

**Supplementary Table S8.** PRISM secondary screen detailing drugs exhibiting significant shifts in sensitivity between Mutated and NonMutated HNSC cell lines.

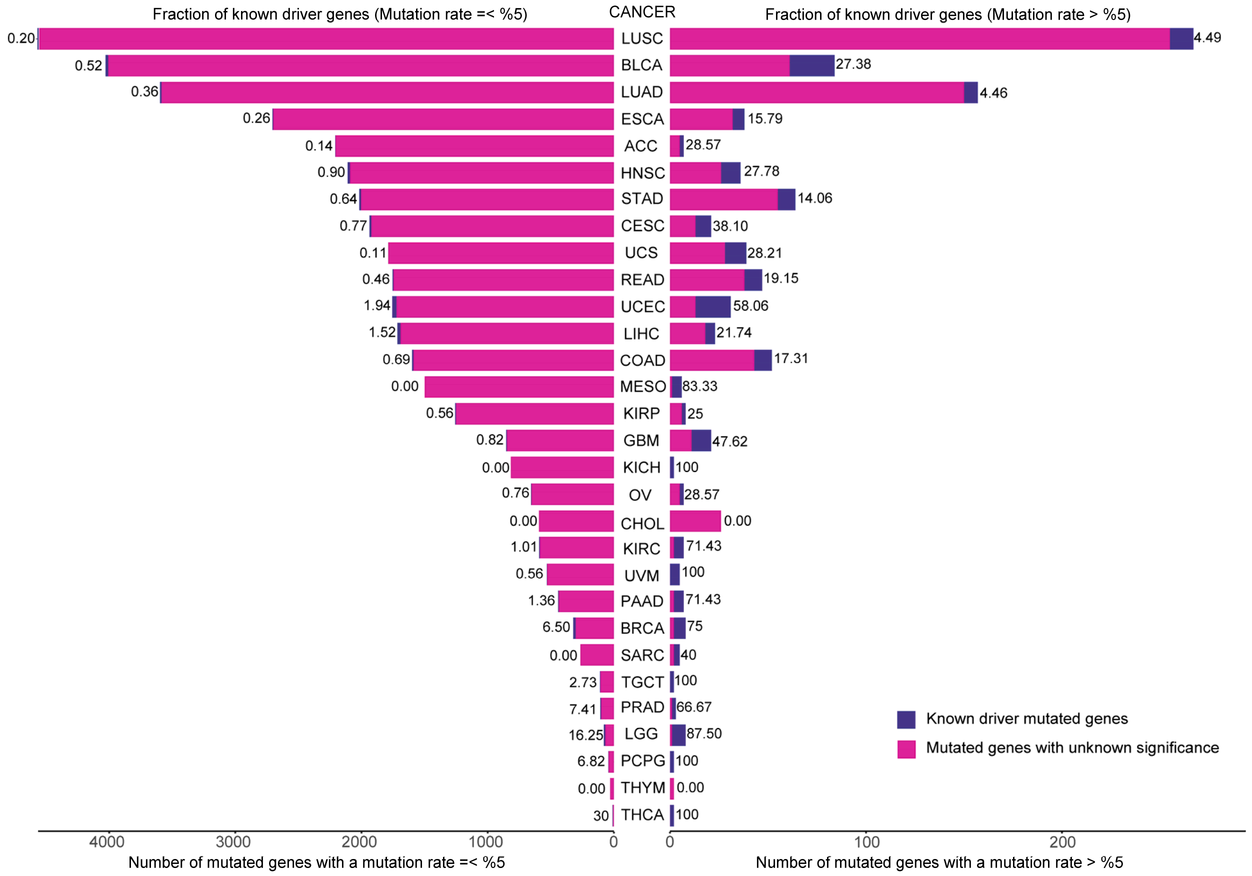


**Supplementary Fig. 1. Fraction of known drivers amongst recurrently mutated genes across diverse cancer types.** Somatic mutations across 30 types of solid tumors were obtained from the Pan-Cancer atlas dataset, comprising ~10K tumor samples. Stacked bar-plot showing the fraction of the previously known driver genes to the unknown mutated genes, where the Y-axis indicating the cancer types, and the X-axis denotes the number of mutated genes in each cancer. Cancer types are sorted from top to bottom based on total number mutated genes. Recurrent mutated genes (mutation rates > %0.5 on the right) depicted a higher fraction of known driver genes, while only a very small fraction of rarely mutated genes (mutation rates < 0.5% on the left) are known yet. LUSC, lung squamous cell carcinoma; BLCA, bladder urothelial carcinoma; LUAD, lung adenocarcinoma; ESCA, oesophageal carcinoma; ACC, adrenocortical carcinoma; HNSC, head and neck squamous carcinoma; STAD, stomach adenocarcinoma; CESC, cervical squamous cell carcinoma and endocervical adenocarcinoma; UCS, uterine carcinosarcoma; READ, rectum adenocarcinoma; UCEC, uterine corpus endometrial carcinoma; LIHC, liver hepatocellular carcinoma; COAD, colon adenocarcinoma; MESO, mesothelioma; KIRP, kidney renal papillary cell carcinoma; GBM, glioblastoma multiforme; KICH, kidney chromophobe; OV, ovarian serous cystadenocarcinoma; CHOL, cholangiocarcinoma; KIRC, kidney renal clear cell carcinoma; UVM, uveal melanoma; PAAD, pancreatic adenocarcinoma; BRCA, breast invasive carcinoma; SARC, sarcoma; TGCT, testicular germ cell tumours; PRAD, prostate adenocarcinoma;

LGG, brain lower grade glioma; PCPG, pheochromocytoma and paraganglioma; THYM, thymoma; THCA, thyroid carcinoma.

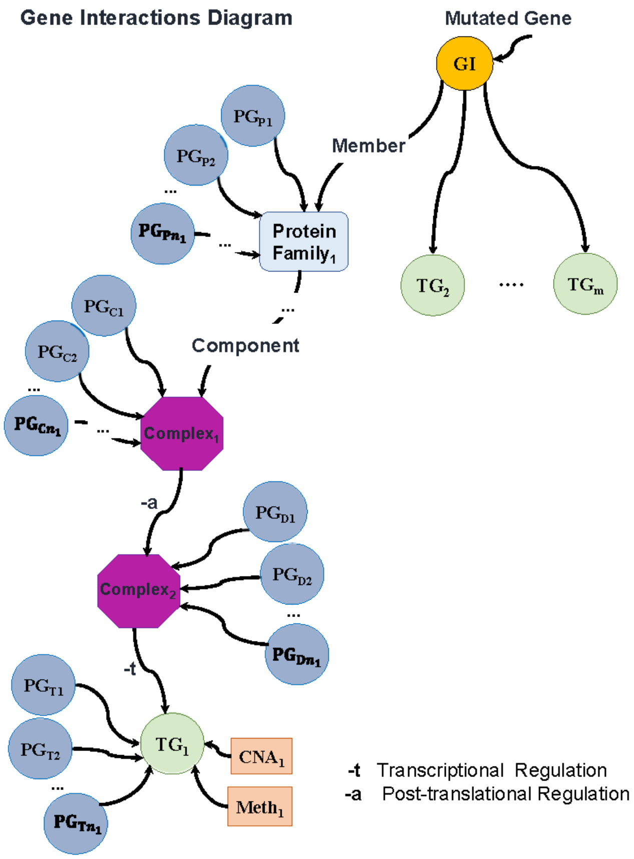


**Supplementary Fig. 2. Schematic illustration of the regulatory interactions of a mutated gene (*GI*) and its downstream transcriptional target genes (*TG*s).** Shown is an example gene regulatory influence network from a mutated gene of interest, *GI,* to its transcriptional target gene, $TG_i$, where nodes include *Gene*, *Complex formation, Abstract and Protein family,* whereas, edges include *Transcriptional regulation (-t), Post-transcriptional activation (-a), Member, and Component*. In the transcriptional influence network, indirect transcriptional TG of a *GI* refers to the TGs, that include only one post-transcriptional activation interaction (-a) in the path of *GI* to *TG* followed by a transcriptional regulation (-t) to *TG.* In contrast, in the path of GI to a direct TG, no post-transcriptional activation interaction (-a) is included. All of *trans*-regulatory factors (*PG*s) in the path of *GI* to *TG* and *cis*-regulatory factors of a *TG* (DNAMeth and sCNA) are considered as a potential confounders by the SYS-Mut model.

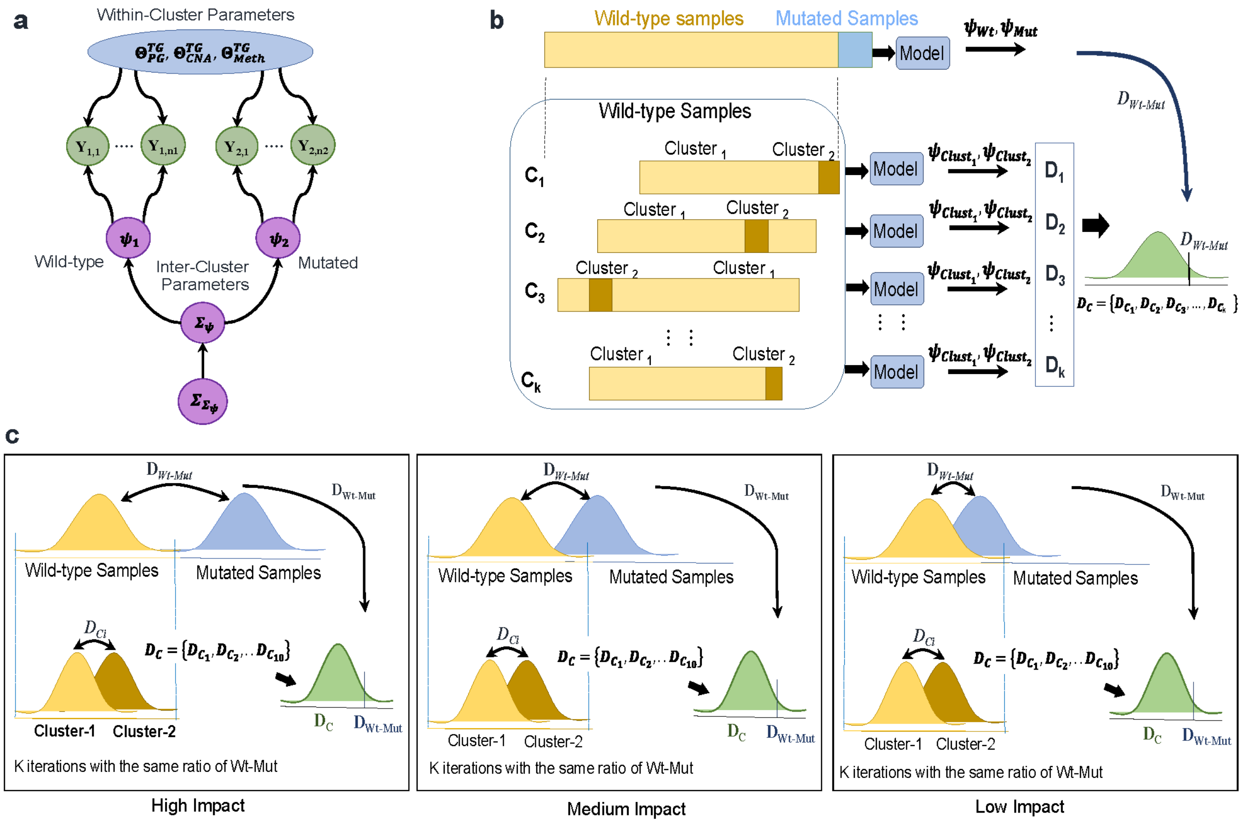


**Supplementary Fig. 3. Detailed methodology to identify mutated genes with significant impact in primary tumor datasets. a** Estimation of the inter- and intra-cluster parameters for $n_1$ Wildtype and $n_2$ Mutated samples. **b** Shown are the quantifications of mutation impact per *TG*. Bhattacharyya distance between the estimated probability distributions of wildtype vs. mutated samples are denoted by ($D_{Wt-Mut}$). The Bhattacharyya distances of '*clean wild-type'* samples, randomly split into two sub-clusters in *K* iterations, are indicated by $\boldsymbol{D}_c = \{D_{c_1}, D_{c_2}, \ldots, D_{c_k}\}$. **c** Depicts High, Medium, and Low level of transcriptional impacts, quantified by comparison of $D_{Wt-Mut}$ and the distribution of $\boldsymbol{D}_c = \{D_{c_1}, D_{c_2}, \ldots, D_{c_k}\}$.

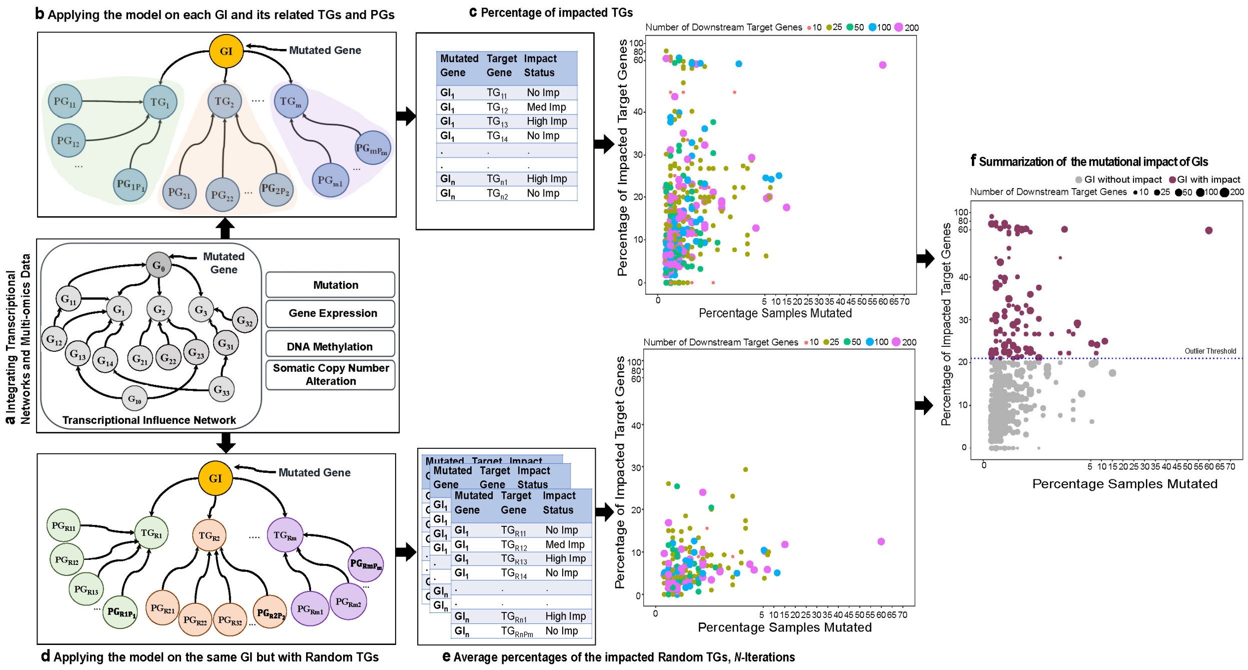


**Supplementary Fig. 4. Network based summarization of mutational impact of *GI* using a network randomization strategy. a** Capturing the *cis*- and *trans*-regulatory confounders effect, for each identified transcriptional target genes (*TG*) of the mutated gene *GI*, by integrating the molecular profiles of the tumor samples including mutation status of the gene of interest, gene expression levels of all genes of the influence network, along with DNA Methylation and somatic copy number alterations (sCNA) of all target genes. **b** Applying the model on each mutated *GI* and its downstream *TG*s and other trans-regulators. **c** Estimation of the percentage of impacted target genes associated with each *GI* (Y-Axis) against of the *GI*'s mutation rate (X-Axis). **d** Applying the model to the same mutated *GI* but randomly substituted *TG*s with their respective *trans*-regulatory factors ($PG_{RS}$). **e** Shown are *N* iterations to estimate impact of a *GI* on its randomly substituted TGs. Colored dots indicate the average fractions of impacted random target genes associated with each *GI* (Y-Axis) against the *GI*'s mutation rate (X-Axis). **f** Scatter plot depicting the comparison of probability scores of the regulatory impact for each *GI* with real transcriptional targets as compared to the average of randomly replaced target genes.

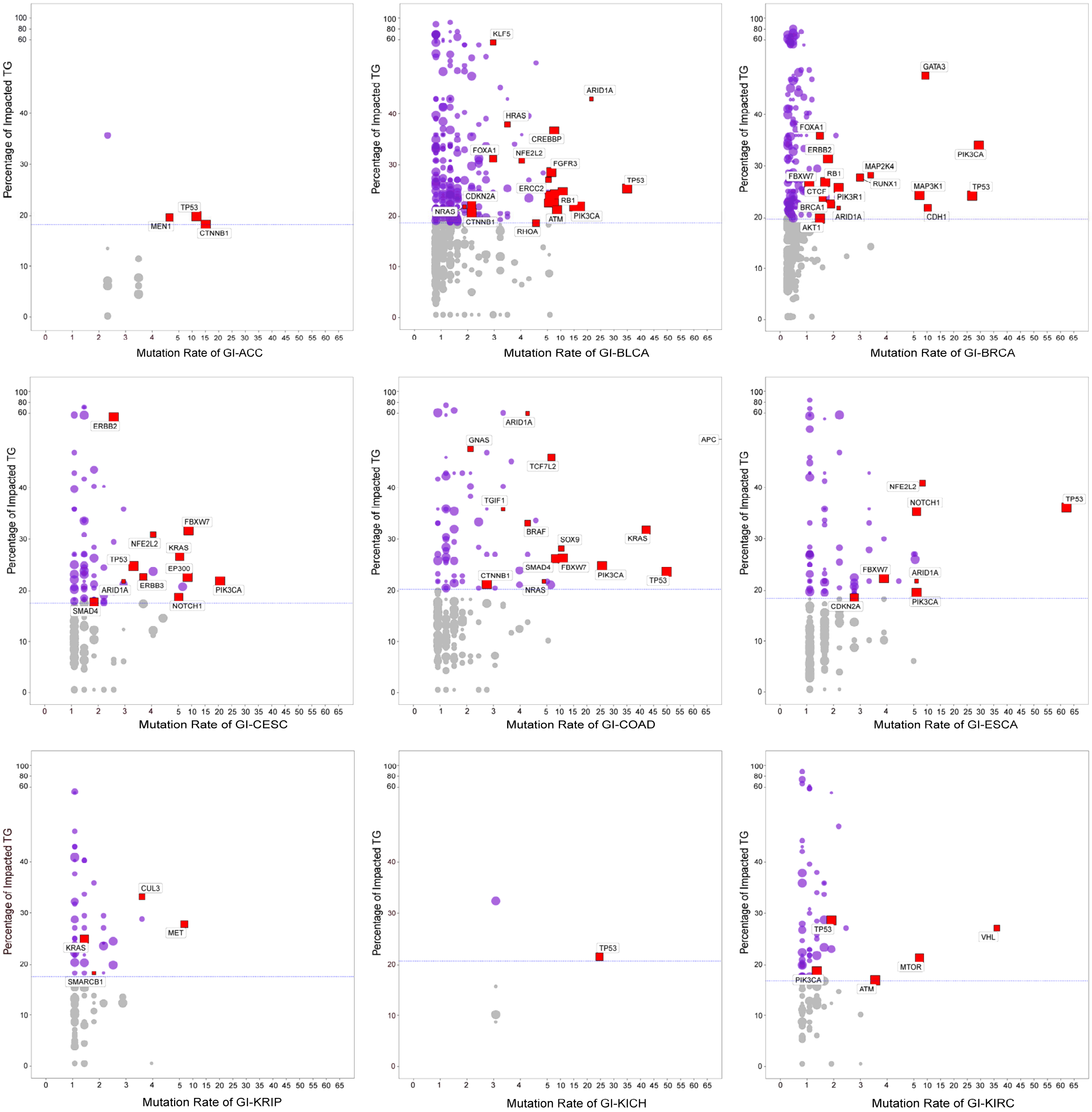
Percentage of Impacted TG
MEN1
TP53
CTNNB1
Mutation Rate of GI-ACC
KLF5
ARID1A
HRAS
CREBBP
FOXA1
NFE2L2
FGFR3
TP53
ERCC2
CDKN2A
RB1
NRAS
ATM
PIK3CA
CTNNB1
RHOA
Mutation Rate of GI-BLCA
GATA3
FOXA1
ERBB2
PIK3CA
MAP2K4
FBXW7
RB1
RUNX1
CTCF
PIK3R1
MAP3K1
TP53
BRCA1
ARID1A
CDH1
AKT1
Mutation Rate of GI-BRCA
ERBB2
FBXW7
NFE2L2
KRAS
TP53
EP300
ERBB3
ARID1A
PIK3CA
SMAD4
NOTCH1
Mutation Rate of GI-CESC
ARID1A
GNAS
APC
TCF7L2
TGIF1
BRAF
SOX9
KRAS
SMAD4
FBXW7
PIK3CA
TP53
CTNNB1
NRAS
Mutation Rate of GI-COAD
NFE2L2
NOTCH1
TP53
FBXW7
ARID1A
CDKN2A
PIK3CA
Mutation Rate of GI-ESCA
CUL3
MET
KRAS
SMARCB1
Mutation Rate of GI-KRIP
TP53
Mutation Rate of GI-KICH
TP53
VHL
MTOR
PIK3CA
ATM
Mutation Rate of GI-KIRC

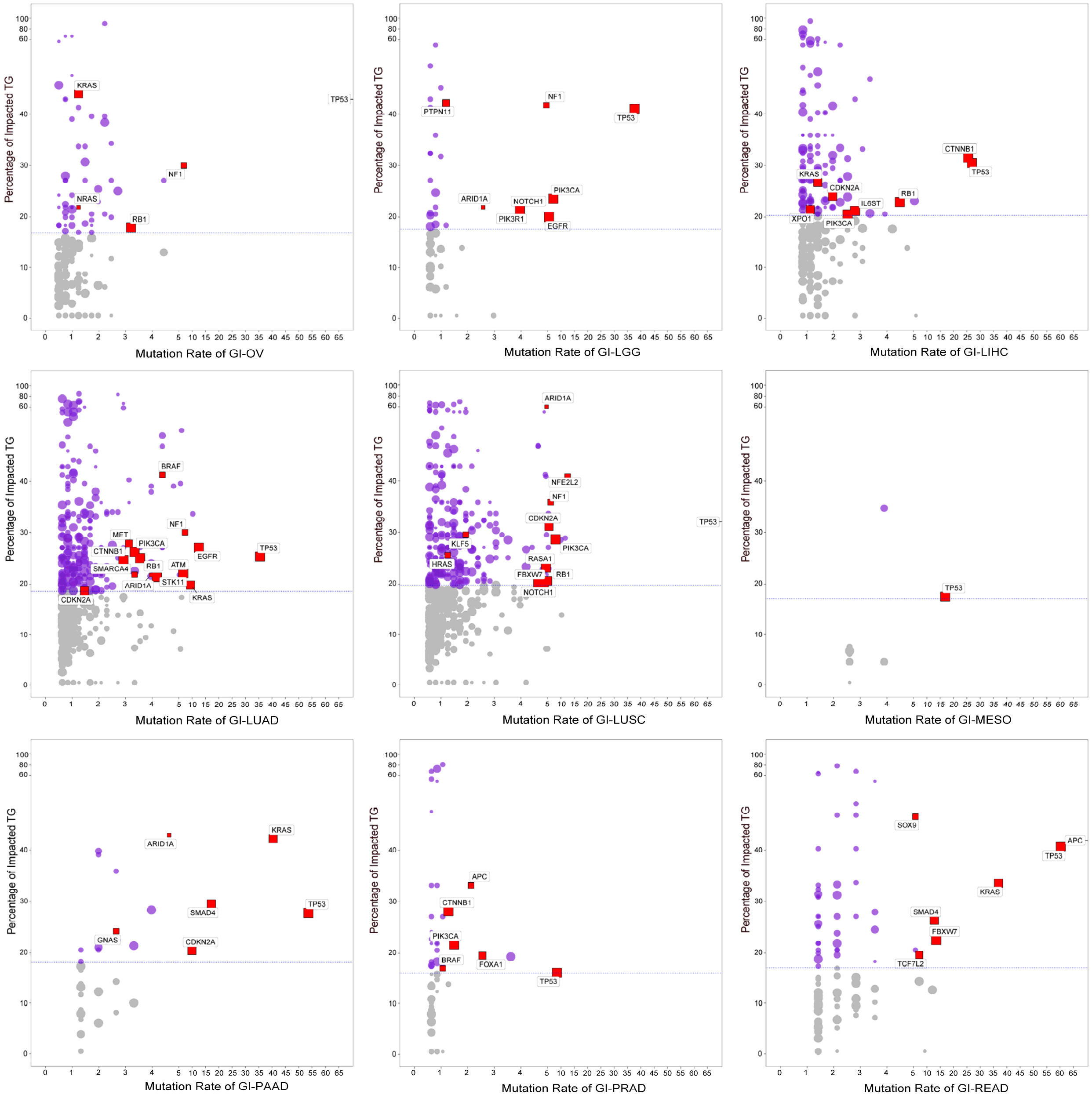
Percentage of Impacted TG
Mutation Rate of GI-OV
KRAS
TP53
NF1
NRAS
RB1
Mutation Rate of GI-LGG
PTPN11
NF1
TP53
ARID1A
NOTCH1
PIK3CA
PIK3R1
EGFR
Mutation Rate of GI-LIHC
CTNNB1
TP53
KRAS
CDKN2A
RB1
IL6ST
XPO1
PIK3CA
Mutation Rate of GI-LUAD
BRAF
NF1
MET
PIK3CA
CTNNB1
TP53
EGFR
SMARCA4
RB1
ATM
ARID1A
STK11
CDKN2A
KRAS
Mutation Rate of GI-LUSC
ARID1A
NFE2L2
NF1
CDKN2A
TP53
PIK3CA
KLF5
RASA1
HRAS
FBXW7
RB1
NOTCH1
Mutation Rate of GI-MESO
TP53
Mutation Rate of GI-PAAD
ARID1A
KRAS
TP53
SMAD4
GNAS
CDKN2A
Mutation Rate of GI-PRAD
APC
CTNNB1
PIK3CA
BRAF
FOXA1
TP53
Mutation Rate of GI-READ
SOX9
APC
TP53
KRAS
SMAD4
FBXW7
TCF7L2

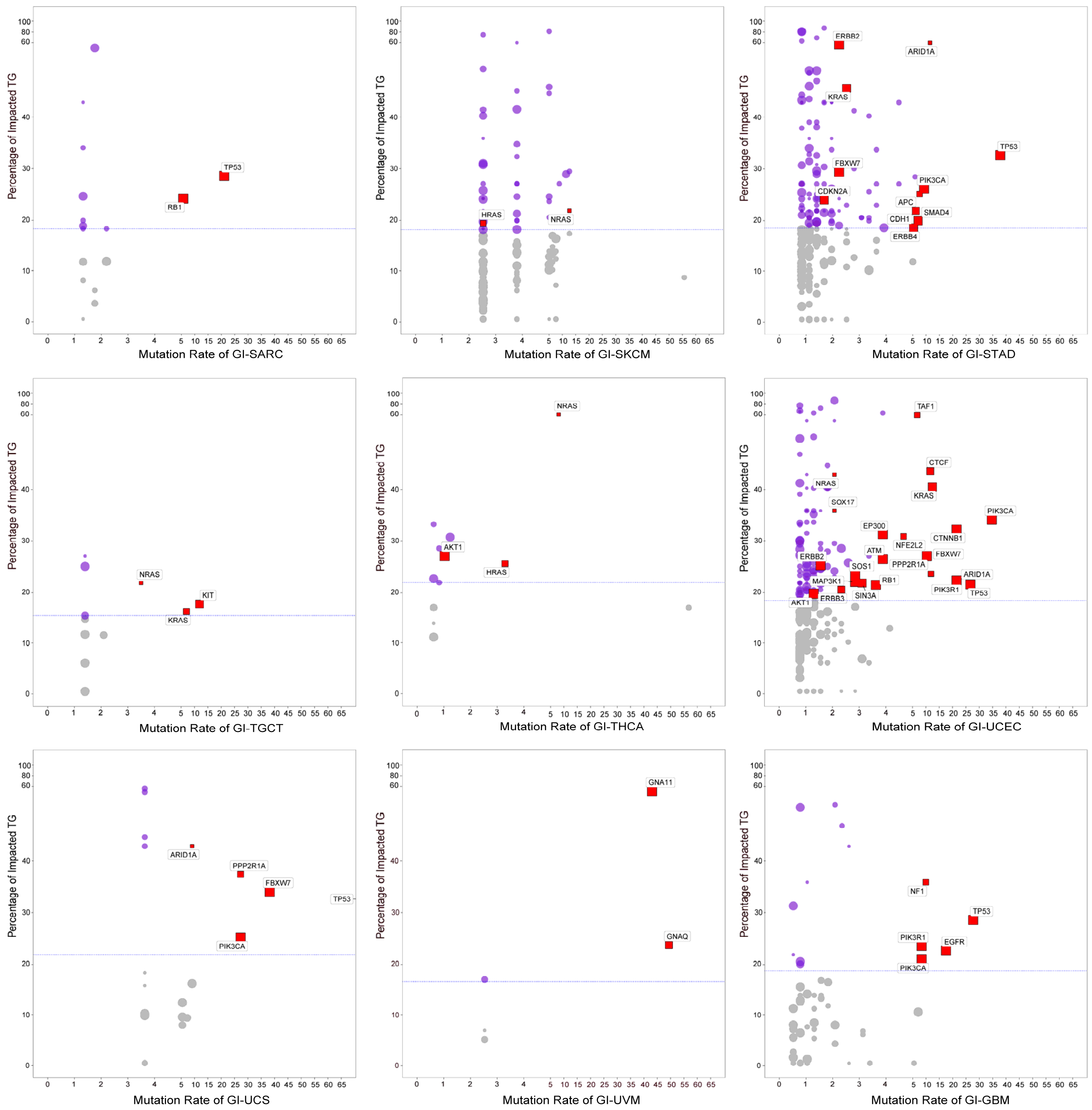
Percentage of Impacted TG
Mutation Rate of GI-SARC
TP53
RB1
Mutation Rate of GI-SKCM
HRAS
NRAS
Mutation Rate of GI-STAD
ERBB2
ARID1A
KRAS
TP53
FBXW7
PIK3CA
CDKN2A
APC
SMAD4
CDH1
ERBB4
Mutation Rate of GI-TGCT
NRAS
KIT
KRAS
Mutation Rate of GI-THCA
NRAS
AKT1
HRAS
Mutation Rate of GI-UCEC
TAF1
CTCF
NRAS
KRAS
SOX17
PIK3CA
EP300
CTNNB1
NFE2L2
ATM
FBXW7
ERBB2
PPP2R1A
SOS1
ARID1A
MAP3K1
RB1
PIK3R1
TP53
AKT1
ERBB3
SIN3A
Mutation Rate of GI-UCS
ARID1A
PPP2R1A
FBXW7
TP53
PIK3CA
Mutation Rate of GI-UVM
GNA11
GNAQ
Mutation Rate of GI-GBM
NF1
TP53
PIK3R1
EGFR
PIK3CA

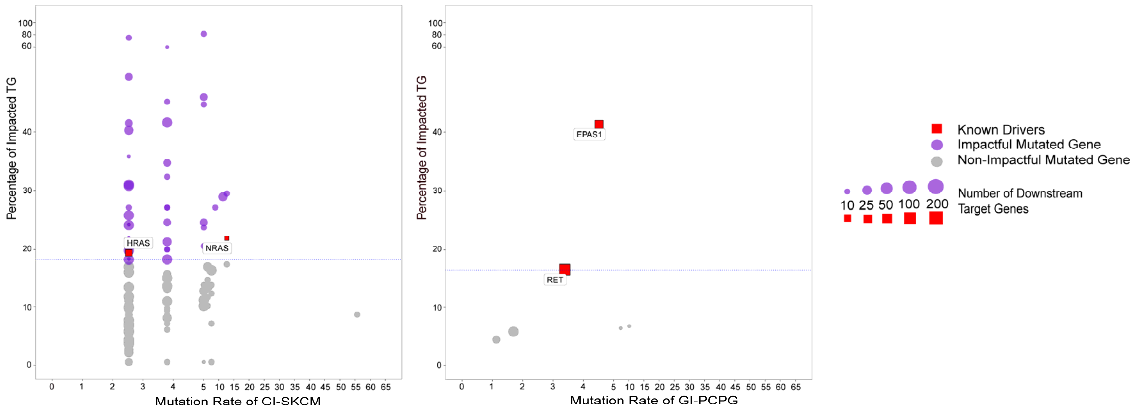


**Supplementary Fig. 5**. **Quantitative analysis of 29 solid tumors of TCGA.** SYS-Mut based quantification of the downstream impact of mutated genes in Pan-Cancer atlas TCGA dataset. Percentage of impacted target genes (Y-Axis) associated with each *GI* is plotted against the *GI*'s mutation rate (X-Axis). The horizontal dashed line denotes the threshold of statistical significance ($P$ < 0.05). The purple and red nodes above the threshold exhibit the all identified mutated genes as significant by SYS-mut, with unknown, and previously known significance, respectively. ACC, adrenocortical carcinoma; BLCA, bladder urothelial carcinoma; BRCA, breast invasive carcinoma; CESC, cervical squamous cell carcinoma and endocervical adenocarcinoma; CHOL, cholangiocarcinoma; COAD, colon adenocarcinoma; ESCA, oesophageal carcinoma; GBM, glioblastoma multiforme; HNSC, head and neck squamous carcinoma; KICH, kidney chromophobe; KIRC, kidney renal clear cell carcinoma; KIRP, kidney renal papillary cell carcinoma; LGG, brain lower grade glioma; LIHC, liver hepatocellular carcinoma; LUAD, lung adenocarcinoma; LUSC, lung squamous cell carcinoma; MESO, mesothelioma; OV, ovarian serous cystadenocarcinoma; PAAD, pancreatic adenocarcinoma; PCPG, pheochromocytoma and paraganglioma; PRAD, prostate adenocarcinoma; READ, rectum adenocarcinoma; SARC, sarcoma; SKCM, skin cutaneous melanoma; STAD, stomach adenocarcinoma; TGCT, testicular germ cell tumors; THCA, thyroid carcinoma; THYM, thymoma; UCEC, uterine corpus endometrial carcinoma; UCS, uterine carcinosarcoma; UVM, uveal melanoma.

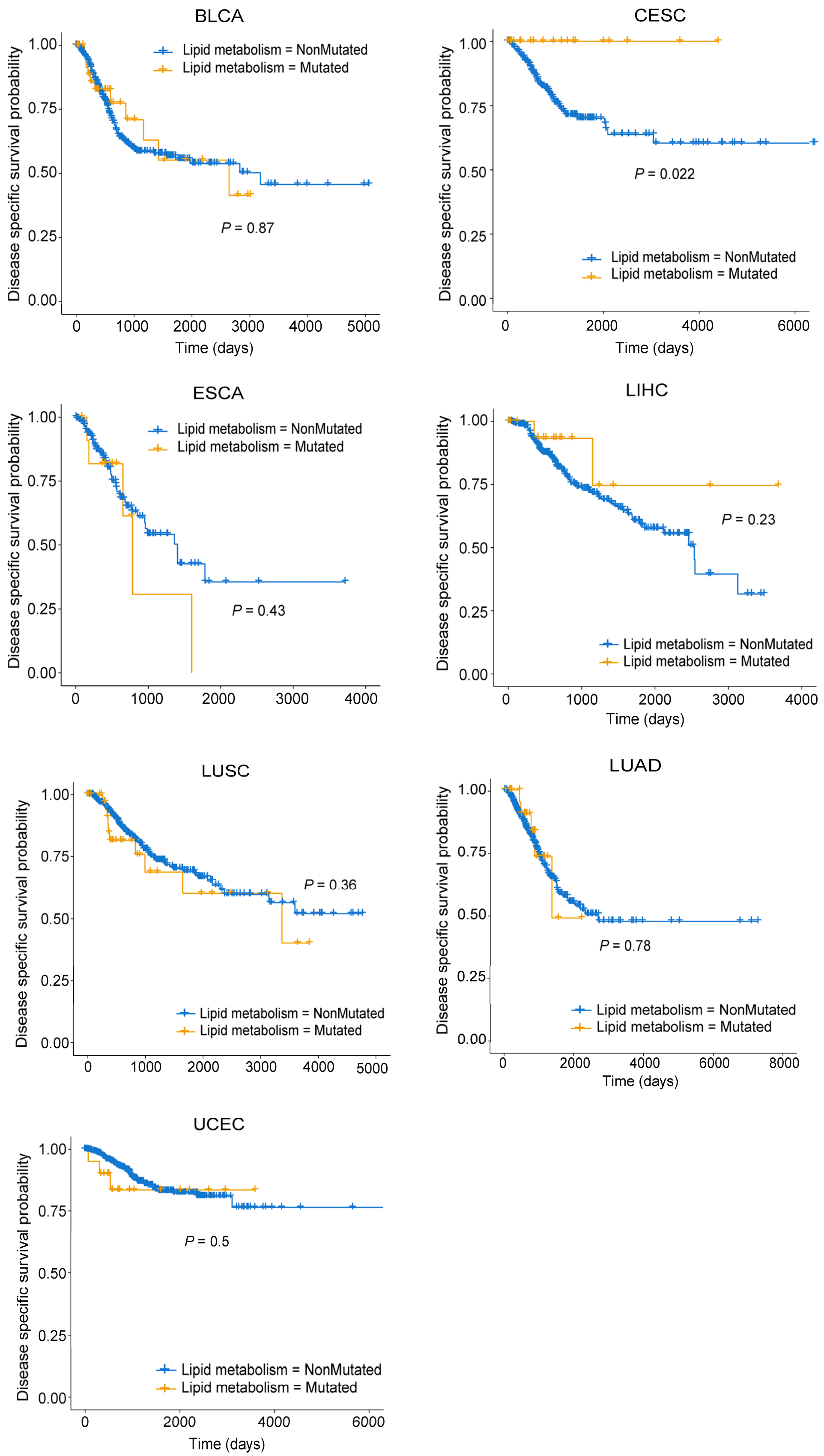
BLCA
Lipid metabolism = NonMutated
Lipid metabolism = Mutated
P = 0.87
Disease specific survival probability
1.00
0.75
0.50
0.25
0.00
0
1000
2000
3000
4000
5000
Time (days)
CESC
Lipid metabolism = NonMutated
Lipid metabolism = Mutated
P = 0.022
0
2000
4000
6000
Time (days)
ESCA
Lipid metabolism = NonMutated
Lipid metabolism = Mutated
P = 0.43
0
1000
2000
3000
4000
LIHC
Lipid metabolism = NonMutated
Lipid metabolism = Mutated
P = 0.23
0
1000
2000
3000
4000
Time (days)
LUSC
P = 0.36
Lipid metabolism = NonMutated
Lipid metabolism = Mutated
0
1000
2000
3000
4000
5000
LUAD
P = 0.78
Lipid metabolism = NonMutated
Lipid metabolism = Mutated
0
2000
4000
6000
8000
Time (days)
UCEC
P = 0.5
Lipid metabolism = NonMutated
Lipid metabolism = Mutated
0
2000
4000
6000
Time (days)

**Supplementary Fig. 6**. **Association of mutations in the lipid metabolism network across cancer types with disease specific survival.** Shown are the Kaplan-Meier estimates of Disease Specific Survival (DSS) for patients with tumors where one or more *GI*s in the lipid metabolism sub-network are Mutated (orange), as compared to their NonMutated (blue) counterparts. The statistical significance of differences in survival rates between Mutated and NonMutated categories was determined using the LogRank test. BLCA, bladder urothelial carcinoma; ESCA, oesophageal carcinoma; LIHC, liver hepatocellular carcinoma; LUAD, lung adenocarcinoma; LUSC, lung squamous cell carcinoma; and UCEC, uterine corpus endometrial carcinoma. To determine the statistical differences of the outcomes, LogRank *P* value are shown.

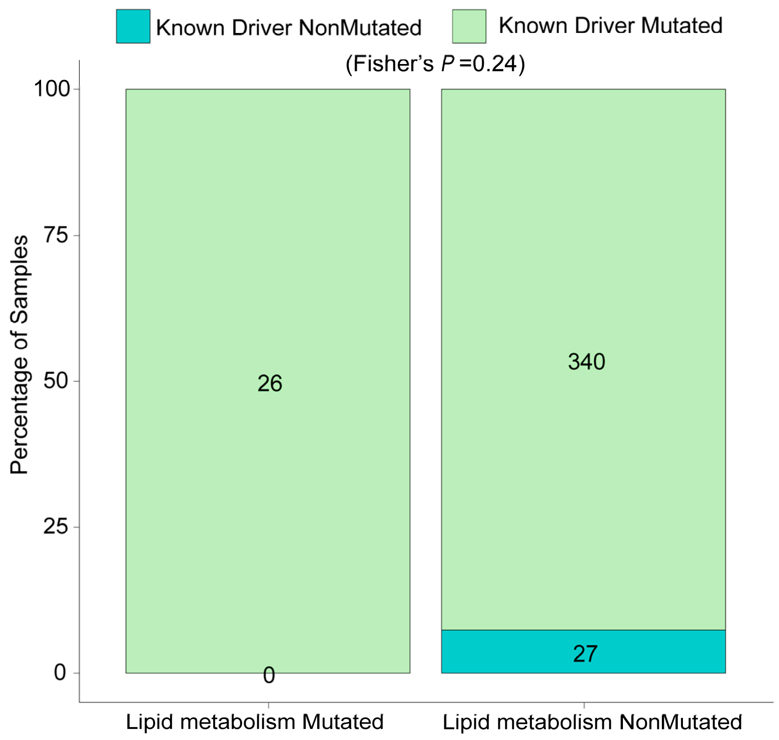


**Supplementary Fig. 7**. **Non-exclusivity of mutations in lipid metabolism network and previously-known HNSC driver genes.** Stacked percentage bar-plot showing the fraction of samples harboring mutations in previously-known HNSC driver genes in either HNSC tumors harboring mutations in the lipid metabolism network (Lipid Metabolism Mutated) or their NonMutated counterparts (Lipid Metabolism NonMutated). Statistical significance was estimated using a Fisher exact test.

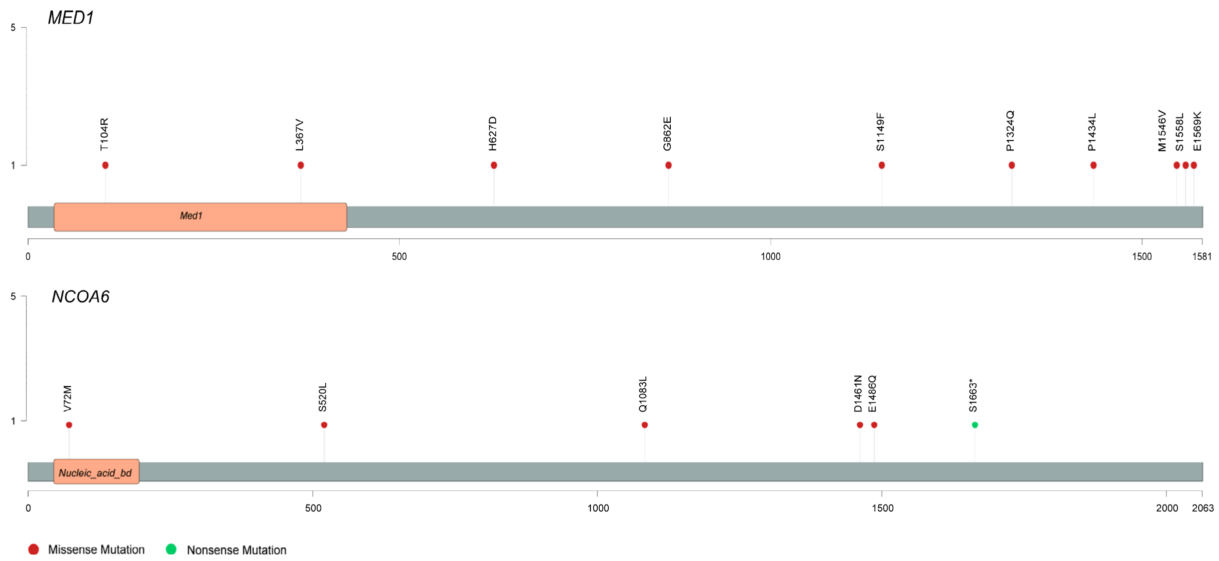


**Supplementary Fig. 8. Mutational hotspot assessments for *MED1* and *NCOA6* in Head and Neck Squamous Cell Carcinoma.** Lolliplot showing somatic mutations in the HNSC dataset across the length of the amino-acid sequences for each of the top mutated genes, *MED1* and *NCOA6* in the lipid metabolism network. Note the broad distribution of mutations throughout the length of the amino-acid sequences for each of the genes in this network, a pattern suggestive of loss-of function as opposed to activating mutations.

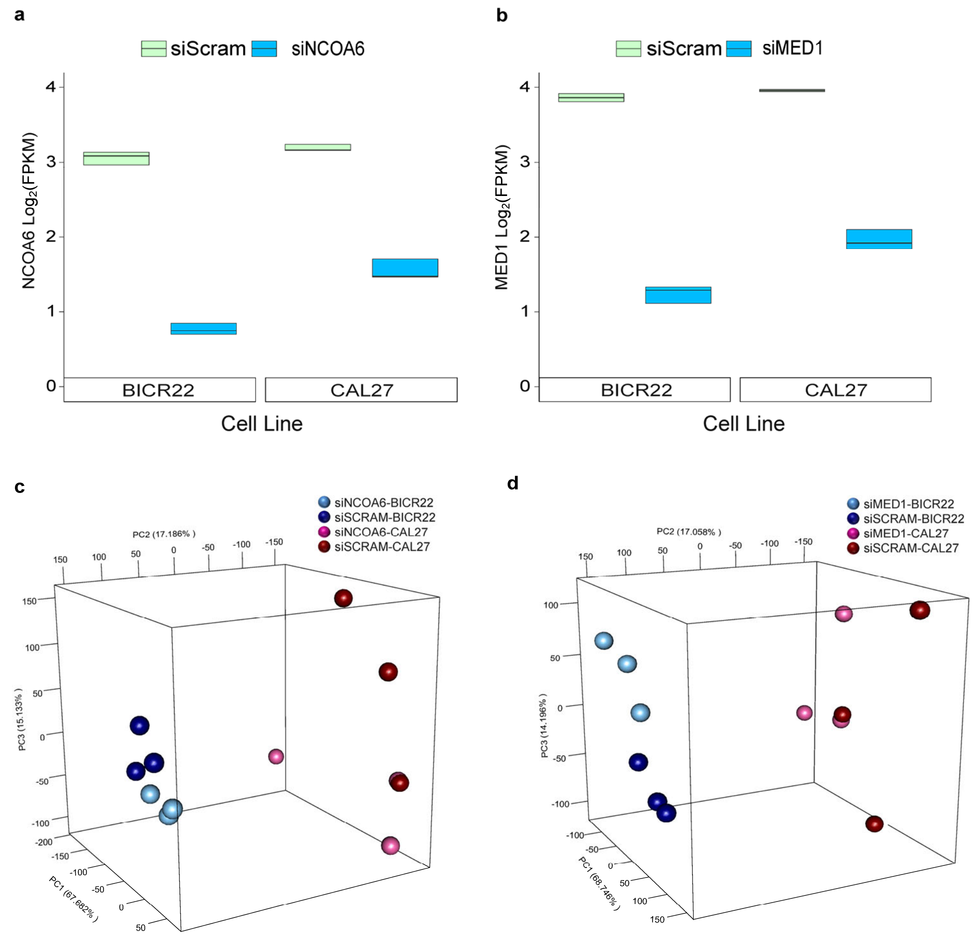


**Supplementary Fig. 9. RNAseq profiling of distinct HNSC cell lines upon siRNA-based knockdown of *MED1* and *NCOA6*. a-b** Boxplots depicting the expression levels (log2FPKM) of *NCOA6* (**a**) and *MED1* (**b**) using RNASeq-based whole transcriptome profiles generated in two distinct HNSC cell lines (BICR22 and CAL27) after 48-hrs of treatment with either siRNA targeting *NCOA6* (siNCOA6) or *MED1* (siMED1), as compared to non-targeting control (siSCRAM). **c-d** Principal component analysis (PCA) of RNASeq-based whole-transcriptome profiles derived from two distinct HNSC cell lines (BICR22 and CAL27) after 48-hrs of treatment with either (**c**) siRNA targeting *NCOA6* (siNCOA6) or (**d**) *MED1* (siMED1), as compared to non-targeting control (siSCRAM). Note the biologic replicates from each cell line and treatment category clustering together.
